# Mendelian imputation of parental genotypes for genome-wide estimation of direct and indirect genetic effects

**DOI:** 10.1101/2020.07.02.185199

**Authors:** Alexander I. Young, Seyed Moeen Nehzati, Chanwook Lee, Stefania Benonisdottir, David Cesarini, Daniel J. Benjamin, Patrick Turley, Augustine Kong

## Abstract

Associations between genotype and phenotype derive from four sources: direct genetic effects, indirect genetic effects from relatives, population stratification, and correlations with other variants affecting the phenotype through assortative mating. Genome-wide association studies (GWAS) of unrelated individuals have limited ability to distinguish the different sources of genotype-phenotype association, confusing interpretation of results and potentially leading to bias when those results are applied – in genetic prediction of traits, for example. With genetic data on families, the randomisation of genetic material during meiosis can be used to distinguish direct genetic effects from other sources of genotype-phenotype association. Genetic data on siblings is the most common form of genetic data on close relatives. We develop a method that takes advantage of identity-by-descent sharing between siblings to impute missing parental genotypes. Compared to no imputation, this increases the effective sample size for estimation of direct genetic effects and indirect parental effects by up to one third and one half respectively. We develop a related method for imputing missing parental genotypes when a parent-offspring pair is observed. We provide the imputation methods in a software package, SNIPar (single nucleotide imputation of parents), that also estimates genome-wide direct and indirect effects of SNPs. We apply this to a sample of 45,826 White British individuals in the UK Biobank who have at least one genotyped first degree relative. We estimate direct and indirect genetic effects for ∼5 million genome-wide SNPs for five traits. We estimate the correlation between direct genetic effects and effects estimated by standard GWAS to be 0.61 (S.E. 0.09) for years of education, 0.68 (S.E. 0.10) for neuroticism, 0.72 (S.E. 0.09) for smoking initiation, 0.87 (S.E. 0.04) for BMI, and 0.96 (S.E. 0.01) for height. These results suggest that GWAS based on unrelated individuals provides an inaccurate picture of direct genetic effects for certain human traits.

## Introduction

Genome-wide association studies (GWAS) have found thousands of associations between genetic variants and human traits^1^ and have enabled the prediction of human traits from genetic data through the use of polygenic scores^2^. GWAS typically estimate the additive effect of an allelic substitution at a single nucleotide polymorphism (SNP) by regression of individuals’ phenotypes onto the number of copies of an allele (genotype) that they carry.

Multiple different phenomena can contribute to the effects estimated by GWAS applied to unrelated individuals^3^, which we refer to as ‘population effects’, since they reflect the overall genotype-phenotype association in the population. The causal effect of inheriting a particular allele, called the direct genetic effect, contributes to population effect estimates. However, indirect genetic effects – effects of genetic variants in one individual that affect the trait of another through the environment – from relatives, such as parents, can also contribute to population effect estimates. Such effects have been shown to be important for educational attainment^4,5^. Both direct and indirect genetic effects are causal effects of alleles, due to their presence in the phenotyped individual and/or relatives. It is also possible for the causal effect of an allele to be magnified by assortative mating with respect to a correlated phenotype, which induces correlations between causal alleles at different genomic locations^3,4^. In addition to causal effects, non-causal associations between genotype and phenotype due to population stratification can contribute to population effect estimates^6,7^. Population stratification occurs when different subpopulations within the GWAS sample have different mean trait values, inducing correlations between SNPs that are differentiated between the subpopulations and the trait.

Decomposing the population effects estimated by GWAS into the different components is important for interpreting and applying GWAS results. For example, polygenic prediction of educational attainment (EA) leverages indirect genetic effects from parents^5,8^. Thus, any potential application of genetic prediction of EA should take account of the fact that a substantial fraction of the predictive ability of the score derives from prediction of the family environment, rather than ‘innate’ abilities of the child. It has also been shown that indirect effects and assortative mating can lead to spurious inference in Mendelian Randomisation, and that this can be remedied by using unbiased estimates of direct genetic effects^9^. Further, subtle population stratification effects in GWAS of height resulted in spurious inference of selection on height in Europe^7,10^, highlighting the need for stratification free estimates of direct genetic effects on traits.

Direct genetic effects can be separated from other sources of genotype-phenotype association by taking advantage of the randomisation of genetic material that occurs during meiosis, which is independent of the environment^8,11–13^. The offspring genotype varies randomly around the expectation given the genotype of the mother and father due to segregation of genetic material in the parents during meiosis. Thus, analysis of parent-offspring trios can be used to estimate direct genetic effects separately from indirect genetic effects and confounding effects^8,14^. However, large samples with genetic data on individuals and both parents are not widely available. A less powerful approach uses genetic differences between siblings, which are also a consequence of random segregations in the parents during meiosis, to estimate direct genetic effects. This approach has been more widely applied due to the greater availability of large samples of genotyped sibling pairs^5,7,15,16^.

Genetic data on siblings contains information about the genotypes of the parents of the siblings. We develop a method for imputing missing parental genotypes from sibling genotypes. This method takes advantage of the fact that, given knowledge of whether siblings inherited the same or different alleles from each parent --- i.e., given knowledge of the identity-by-descent (IBD) states of the siblings’ alleles --- the parental alleles that have been observed in the siblings can be determined. Unlike methods based on differences between sibling genotypes, our method can provide unbiased estimates of indirect genetic effects from siblings and distinguish them from direct effects and indirect genetic effects from parents.

We also show that imputing the missing parent’s genotype when genotypes are available for a phenotyped individual (proband) and one of the proband’s parents enables unbiased estimation of direct genetic effects. We provide the methods for imputing missing parental genotypes in a software package, SNIPar (single nucleotide imputation of parents), that can also infer genome-wide direct and indirect effects of SNPs. The software package models phenotypic correlations within-families and can also be applied to samples of probands with both parents genotyped.

We apply our methods to 45,826 White British individuals in the UK Biobank with at least one genotyped parent or sibling. We estimated direct and indirect genetic effects for ∼5 million genome-wide SNPs for educational attainment, height, body mass index (BMI), neuroticism, and smoking initiation. We use the resulting direct effect estimates to estimate the genetic correlation between direct genetic effects and population effects for each of the traits. Our findings suggest that for certain traits, SNP effects estimated from standard GWAS provide inaccurate estimates of direct genetic effects.

## Results

### Imputing parental genotypes from sibling genotypes

Given genotype observations at a single SNP for a sibling pair, the number of parental alleles that have been observed depends upon the whether the siblings have inherited the same or different alleles from each parent, i.e. the IBD states of the alleles (Figure 1).

**Figure 1.**
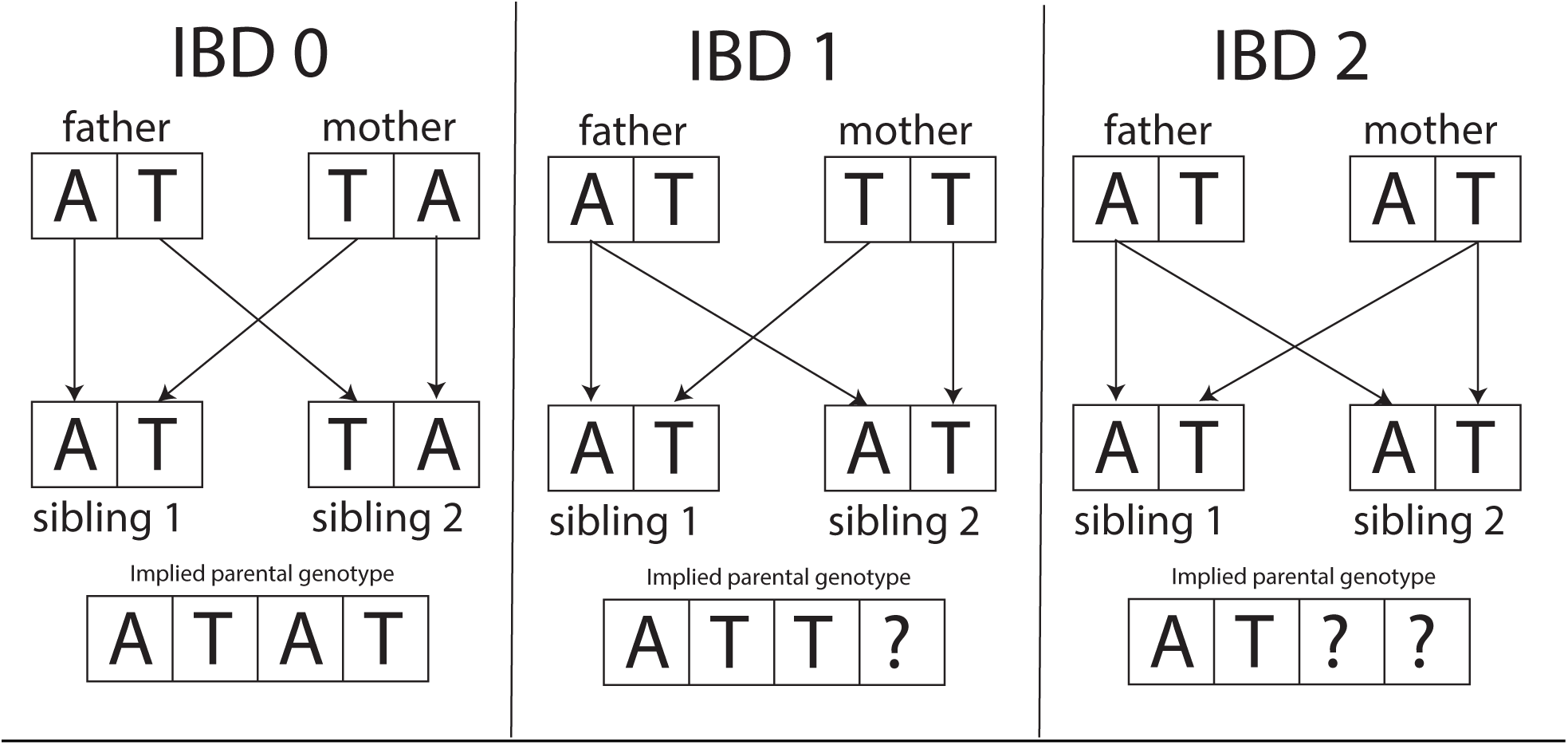
Imputation of parental genotype from sibling genotype. Here we illustrate how, given knowledge of the IBD state between the siblings’ alleles, the combined maternal and paternal genotype of the parents of the siblings can be imputed. If the siblings do not share any alleles identical-by-descent (IBD), then all four parental alleles are observed (IBD 0). If the siblings share one allele by descent from their parents, then three parental alleles are observed, and one allele is unobserved (IBD 1). If the siblings share both alleles by descent from their parents, then only two parental alleles are observed and two are unobserved (IBD 2).

Since parent-of-origin of alleles cannot be determined from sibling data alone, we impute the sum of maternal and paternal genotypes. Let *g*_par(i)_ = *g*_m(i)_ + *g*_p(i)_ be the sum of the genotype of the mother (*g*_m(i)_) and the genotype of the father (*g*_p(i)_) in family *i*, and let *g*_i1_ and *g*_i2_ be the genotypes of the two siblings. We compute E[*g*_par(i)_|*g*_i1_, *g*_i2_, IBD_i_], where IBD_*i*_ is the IBD state of the two siblings. Since all four alleles are observed in IBD state 0, we have that

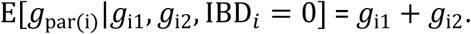

When we do not observe a parental allele, we impute it using the population allele frequency, *f*. Therefore,

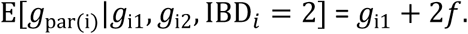

When the siblings share one allele by descent from their parents, it is necessary to know which allele is shared to use all of information in the siblings’ genotypes. Let 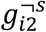 be the binary genotype (presence/non-presence) of the allele in sibling 2 that is not shared IBD with sibling 1. We therefore have

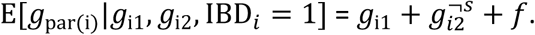

Without phased data, it is impossible to determine which allele is shared when both siblings are heterozygous and in IBD state 1. However, it can be shown that with un-phased data (Supplementary Note)

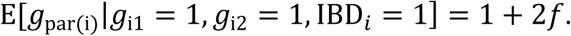

It is therefore possible to perform the imputation without access to phased genotypes, but there is a loss of information compared to imputation with phased genotypes when both siblings are heterozygous and in IBD state 1 (Supplementary Note). We generalise the imputation procedure to families with more than two observed sibling genotypes in the Supplementary Note.

By using the population allele frequencies to impute the unobserved parental alleles, we are assuming that parental alleles are uncorrelated. Apart from in samples exhibiting very strong structure or inbreeding, the correlations between parental alleles at individual SNPs will be very weak, so this assumption will be approximately correct.

### Estimating effects using sibling and imputed parental genotypes

We consider a model for the effect of a SNP on the traits of two siblings that includes both direct genetic effects and indirect genetic effects from parents and siblings. Let *Y*_ij_ be the phenotype of sibling *j* in family *i*. Then

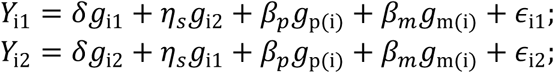

where *δ* is the direct effect of the SNP and *η*_*s*_ is the indirect genetic effect from the sibling. Offspring genotypes are conditionally independent of environmental effects given parental genotypes. Therefore, estimates of direct effects and indirect genetic effects from fitting this model are unbiased (Supplementary Note)^11^. Any correlation between the siblings’ genotypes and other factors affecting the trait are captured by the parental genotypes. Thus, *β*_*p*_ and *β*_*m*_ capture indirect genetic effects from the father and mother respectively, in addition to any confounding due to population stratification and any magnification of direct and sibling indirect genetic effects due to assortative mating^11^ (Supplementary Note). We will refer to *β*_*p*_ and *β*_*m*_ as ‘parental effects’, even though they can reflect phenomena other than indirect genetic effects from parents. Further, the residuals *ϵ*_i1_ and *ϵ*_i2_ are uncorrelated with the genotypes of the siblings and parents, but may be correlated with each other (Supplementary Note). Note that standard GWAS methods that regress proband phenotype onto proband genotype are expected to estimate *δ* + *η*_*s*_/2 + (*β*_*p*_ + *β*_*m*_)*p*2, which we refer to as ‘population effects’.

Many previous analyses of family data regressed the difference in sibling phenotype onto the difference in sibling genotype^5,7,16^. In our model, this corresponds to:

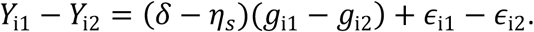

This method is expected to yield unbiased estimates of *δ* − *η*_*s*_ (see our companion paper for further details^17^). The difference between sibling phenotypes forms one axis of information, but there is an orthogonal axis of information: the sum of the sibling phenotypes, which is uncorrelated with the difference between sibling phenotypes. In our model, this corresponds to:

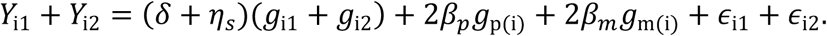

However, for this axis, regression on observed sibling genotypes alone cannot separate direct genetic effects from parental effects. To separate direct genetic effects from parental effects, the imputed parental genotypes derived above can be used. Let ĝ_par(i)_ be the parental genotype imputed from sibling genotypes and IBD information, then, by performing the regression

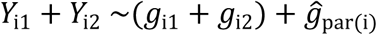

one obtains estimates of *δ* + *η*_*s*_ and the combined parental effect (*β*_*p*_ + *β*_*m*_). These estimates can then be combined with the estimate of *δ* − *η*_*s*_ from differences between siblings to produce separate estimates of *δ* and *η*_*s*_. We note that the regressions outlined above do not generalise well to samples with families with different numbers of siblings and missing phenotype observations. For applications to real data and in our software package, we use a more flexible linear mixed model approach that models correlations between siblings’ phenotypes (Methods and Supplementary Note). Further, we prove that the method we propose gives estimates of the effects that converge to the true values (Supplementary Note).

While our method is able to distinguish indirect genetic effects from siblings from direct genetic effects, more precise estimates of direct genetic effects can be obtained by assuming that *η*_*s*_ = *m* (Supplementary Note), at the cost of some bias if *η*_*s*_ ≠ *m* (see our companion paper for details^17^). For the following results, we make the assumption that *η*_*s*_ = *m* unless otherwise stated.

By using imputed parental genotypes, more precise estimates of *δ* can be produced than from using sibling differences alone: in the Supplementary Note, we show that by using imputed parental genotypes derived from phased IBD data, the effective sample size for estimation of *δ* is increased by a factor of 2(2 + *r*)*/*(3 + 3*r*), where r is the correlation of the siblings’ residuals, which will be approximately equal to their phenotypic correlation for polygenic traits. For *r* > 0, this has a maximum of 4/3 at *r* = 0, corresponding to an effective sample size gain of 1/3^rd^ (Figure 2).

**Figure 2.**
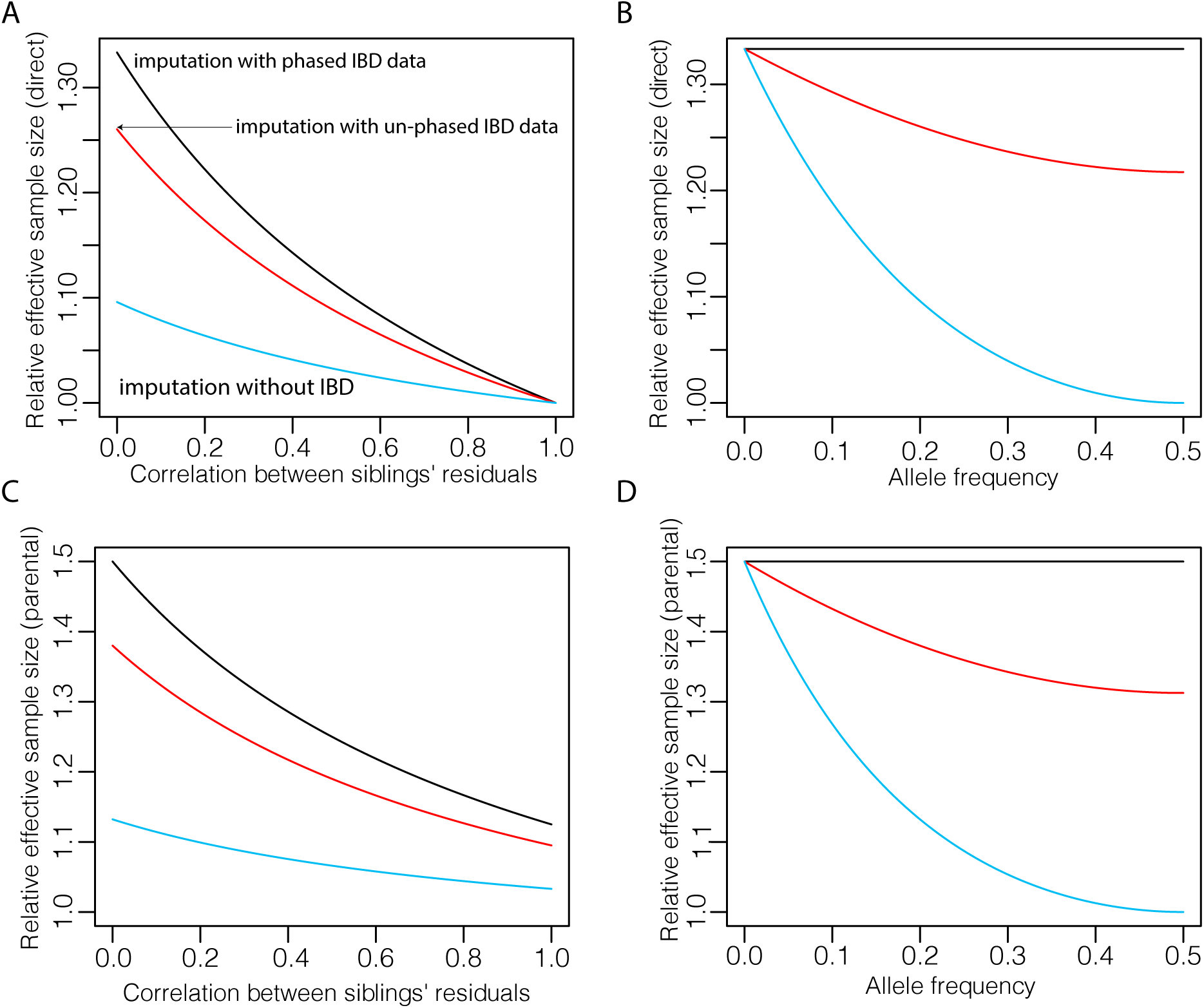
Relative efficiency for estimation of direct and parental effects using different imputation methods. We compare the theoretical effective sample size for estimation of direct genetic effects and combined parental effects from three imputation methods: one that does not use identity-by-descent (IBD) segments (blue)^18^, one that uses un-phased IBD segments (red), and one that uses phased IBD segments (black). Effective sample size is measured relative to that from using sibling genotypes alone without any imputation and assuming that we have a sample of independent families with two genotyped and phenotyped siblings in each family (Supplementary Note). **A**) Effective sample size for estimation of the direct genetic effect of an allele with frequency 20% as a function of correlation between siblings’ residuals, which is approximately equal to phenotypic correlation for polygenic traits. **B**) Effective sample size for estimation of direct genetic effects as a function of allele frequency when the correlation between siblings’ residuals is zero. (Results follow a similar pattern for other sibling correlations.) For imputation with un-phased IBD, when both siblings are heterozygous and share one allele IBD, exactly which parental alleles have been observed cannot be determined (Figure 1), so the imputation averages over the two possibilities. When imputing without using IBD segments at all, uncertainty in inference of observed parental alleles increases rapidly with allele frequency. When phased IBD information is used, the parental alleles that have been observed can always be determined, so the relative efficiency does not depend upon allele frequency. **C**) The same as A but for average parental effects. **D**) The same as for B but for average parental effects.

For estimation of parental effects, using parental genotypes imputed from sibling genotypes and phased IBD data increases effective sample size by a factor of 3(2+r)/(4+4r) compared to using sibling genotypes alone. For estimation of both direct and parental effects, the gain is somewhat lower when un-phased identity-by-descent data is used, depending upon allele frequency and *r* (Figure 2 and Supplementary Note).

### Imputing a missing parental genotype from a parent-offspring pair

Consider imputing the genotype of a father whose genotype is unobserved given observations of the proband and the mother’s genotype. More formally, we impute the missing paternal genotype as the expectation given the proband and mother’s genotype: *ĝ*_p(i)_= E[*g*_p(i)_|*g*_*i*1_, *g*_m(i)_]. It is trivial to infer which allele was inherited from the father except when both proband and mother are heterozygous. For example, if the parental genotype is AA and the proband genotype is AT, then the T allele must have been inherited from the missing parent. This means that one half of the paternal genotype can be inferred exactly, and the expectation of the other half is given by the population allele frequency. When both mother and proband are heterozygous, E[*g*_p(i)_|*g*_*i*1_, *g*_m(i)_] can be computed by averaging over the two possible inheritance patterns (Table 1 and Supplementary Note).

**Table 1.** The imputed parental genotype as a function of the proband and maternal genotype: *ĝ*_*p(i)*_= E[*g*_*p(i)*_ |*g*_*i*1_, *g*_*m(i)*_] *for an allele with frequency f*

| | | $g_{m(i)}$ | | |
| --- | --- | --- | --- | --- |
|  |  | 0 | 1 | 2 |
| $g_{i1}$ | 0 | $f$ | $f$ | - |
| | 1 | $1+f$ | $2f$ | $f$ |
| | 2 | - | $1+f$ | $1+f$ |

This approach is generalised to the case when multiple full-sibling offspring of the observed parent are genotyped in the Supplementary Note. We note that more sophisticated methods for phasing and determination of parent-of-origin could be applied to improve the imputation in the case when both parent and proband are heterozygous. However, in this paper, we consider this simple imputation method as it is easy to apply to un-phased genotype data on parent-offspring pairs for very large numbers of SNPs.

### Estimating direct and indirect genetic effects using proband and (imputed) parental genotypes

Consider a sample of families where the genotype of the proband and its mother have been observed but the father’s genotype is unobserved. We show that performing the regression

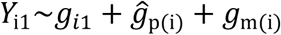

produces estimates of *δ, β*_*p*_, and *β*_*m*_ that converge to their true values (Supplementary Note). This is a consequence of a general proof that we provide for the consistency of least squares estimates of regression coefficients when unobserved covariates are imputed as the conditional expectation given the observed covariates.

The effective sample size for estimation of direct genetic effects relative to complete observation of parental genotypes increases from a minimum of 1/6 when minor allele frequency (MAF) is 0.5 to a maximum of 0.5 as MAF approaches 0 (Supplementary Note and Supplementary Figure 1). The loss in precision compared to complete information is greater when the heterozygosity is higher due to an inability to infer which allele has been inherited from the missing parent when both parent and proband are heterozygous.

### Imputing missing parental genotypes in UK Biobank

We applied our methods to the subsample of the UK Biobank with predominantly white British ancestry^19^. Using KING^20^, we identified 17,296 families with at least two genotyped full siblings but no parents genotyped, giving a total of 19,329 sibling pairs. We inferred un-phased IBD segments for these sibling pairs using KING (Methods). We chose to use un-phased IBD segments as the computational costs of inferring phase for millions of SNPs are high.

We validated the IBD inference using 31 families with two siblings and both parents genotyped, where the IBD state can be determined exactly except when both siblings are heterozygous, finding that the IBD state at 98.4% of SNPs was called correctly (Methods and Supplementary Table 1). We imputed the missing parental genotypes using the IBD segments produced by KING and the procedure outlined above for ∼5 million SNPs with INFO>0.99 and MAF>1% (Methods). We validated that the imputed parental genotypes were approximately unbiased estimates of parental genotypes by comparing imputed to observed genotypes in 31 families with two siblings and both parents genotyped, where the imputation was performed ignoring the observed parental genotypes (Methods).

We identified a further 4,418 families with one parent and at least one offspring genotyped, and imputed the missing parental genotypes for each family. We validated that the imputations were approximately unbiased by comparing observed to imputed parental genotypes for 893 families with both parents genotyped, where the imputation was performed ignoring one of the observed parents’ genotypes (Methods).

### Simulations in UK Biobank

We simulated traits for the 17,296 families in the UK Biobank with at least two genotyped siblings but no genotyped parents. We chose 10,000 SNPs randomly from the SNPs with INFO>0.99 and MAF>1% to use as causal SNPs. Let 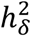 be the proportion of phenotypic variance explained by direct genetic effects. We simulated a trait affected by only direct genetic effects and noise, with 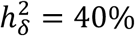 (Methods).

Let 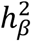 be the proportion of phenotypic variance due to parental effects. For a trait affected by direct and parental effects, the fraction of phenotypic variance explained by population effects, 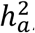, depends on 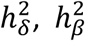, and *r*_δ β_, the correlation between direct and parental effects: 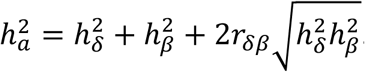. This is because population effects are the sum of direct and average parental effects.

We simulated 40 independent replicates of a trait affected by both direct genetic effects and parental effects (Methods), with 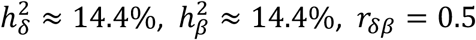, so that 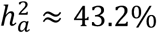. Since there are only a small number of families where both parents are genotyped in the UK Biobank, we used the imputed parental genotypes in place of observed parental genotypes for this simulation. In expectation, this should give the same estimated parental effects as using observed parental genotypes (Supplementary Note). The correlation between direct and population effects, *r*_δ*a*_, is equal to

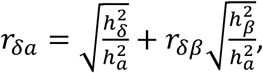

implying a correlation of 0.866 between direct and population effects for the simulated trait.

We estimated direct and average parental effects by regressing the phenotype onto the proband and imputed parental genotype using our mixed model approach to model phenotypic correlations between siblings (Methods). We did this for each trait replicate for ∼5 million SNPs with INFO>0.99 and MAF>1%. We confirmed that direct effect estimates were approximately unbiased for the trait affected by direct genetic effects alone, obtaining a regression coefficient of 0.988 (S.E. 0.015) for regression of direct effect estimates onto direct effects. For the trait affected by both direct and parental effects, we obtained a regression coefficient of 0.994 (S.E. 0.003) from regressing direct effect estimates onto direct effects; and a regression coefficient of 1.001 (S.E. 0.003) from regressing parental effect estimates onto parental effects. These results indicate that our method produces approximately unbiased estimates of direct and parental effects for each SNP.

For the trait affected by both direct and parental effects, we used LD-score regression (LDSC)^21^ to estimate the genetic correlation between direct and parental effects and between direct and population effects (Methods). Across the 40 replicates, our average estimate of the genetic correlation between direct and parental effects was 0.497 (S.E. 0.028), very close to the true value of 0.5; the average estimate of the genetic correlation between direct and population effects was 0.888 (S.E. 0.004). These results indicate that LD-score regression can give approximately unbiased estimates of the genetic correlation between direct and parental effects, and between direct and population effects. This is despite the fact that estimates of the variance explained by direct and parental effects showed bias (Supplementary Table 2). This is similar to other reports indicating that LDSC produces more reliable estimates of genetic correlations than of variance components^22^.

### Genome-wide estimation of direct and indirect genetic effects for five traits

We used a sample of 45,826 White British individuals with at least one genotyped sibling or parent, where missing parental genotypes were imputed, to estimate direct and indirect (sibling, paternal, maternal, and average parental) genetic effects of ∼5 million SNPs (MAF>1%) on height, BMI, educational attainment (years), neuroticism score, and whether an individual has ever smoked (“ever-smoked”) (Methods). Traits were adjusted for 40 genetic principal components before SNP effects were estimated.

From the genome-wide estimates of SNP effects, we did not find evidence for a substantial contribution from indirect genetic effects from siblings (Supplementary Table 3), but power for this analysis was limited. In our companion paper, we found no evidence for substantial indirect genetic effects from siblings using a more powerful analysis of polygenic scores^17^. Therefore, in order to increase precision of estimates of direct and parental effects, we estimated effects assuming that indirect genetic effects from siblings were zero.

At these sample sizes, power is limited for analysis of direct and indirect genetic effects of individual SNPs. We therefore focused on estimating the genome-wide correlation between direct genetic effects and population effects using LD-score regression (LDSC) (Methods). This measures the degree to which population effect estimates are biased by indirect genetic effects and population stratification. This correlation reflects the relative amount of signal in population effects coming from direct genetic effects and the correlation between direct and average parental effects. The correlation can also be expressed in terms of the ratio 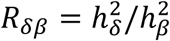:

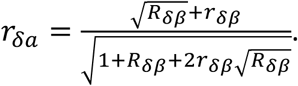

We plot *r*_δ*a*_ as a function of *R*_δ β_ for various values of *r*_δ β_ in Figure 4. This shows that, in order for the correlation between direct and GWAS effects to be substantially below one, the correlation between direct and parental effects must be substantially below one, and that the phenotypic variance explained by direct effects cannot be many times larger than the variance explained by parental effects.

The estimated genetic correlation between direct and population effects ranged from 0.61 (S.E. 0.093) for EA to 0.96 (S.E. 0.015) for height (Supplementary Table 4 and Figure 3). All of the genetic correlation estimates were statistically significantly below 1 (P<0.005, one-sided Z-test). We also estimated genetic correlations between direct and parental effects, but these estimates had low precision (Supplementary Table 4). We obtained consistent results from estimates of the genetic correlation between direct effect estimates and publicly available GWAS summary statistics (Supplementary Table 5).

**Figure 3.**
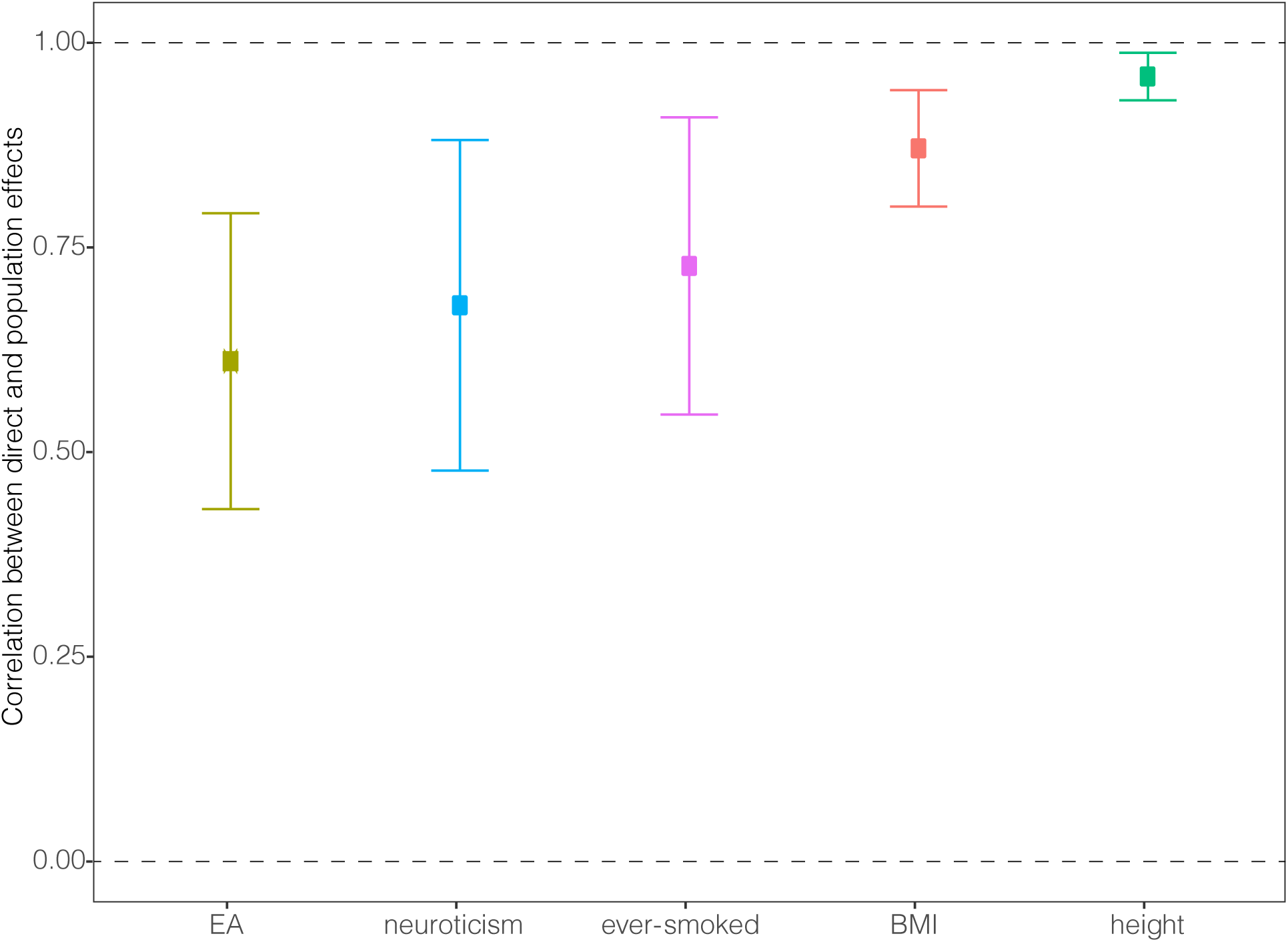
Estimates of genetic correlation between direct effects and population effects. The estimate is given along with the 95% confidence interval. Population effects include direct effects, indirect effects from relatives, magnification due to assortative mating, and bias due to population stratification. Estimates were derived from applying LD-score regression to direct and population effect estimates, derived from a sample of White British individuals from the UK Biobank, for ∼5 million SNPs with MAF>1% (Methods). Traits were adjusted for 40 genetic principal components prior to analysis.

**Figure 4.**
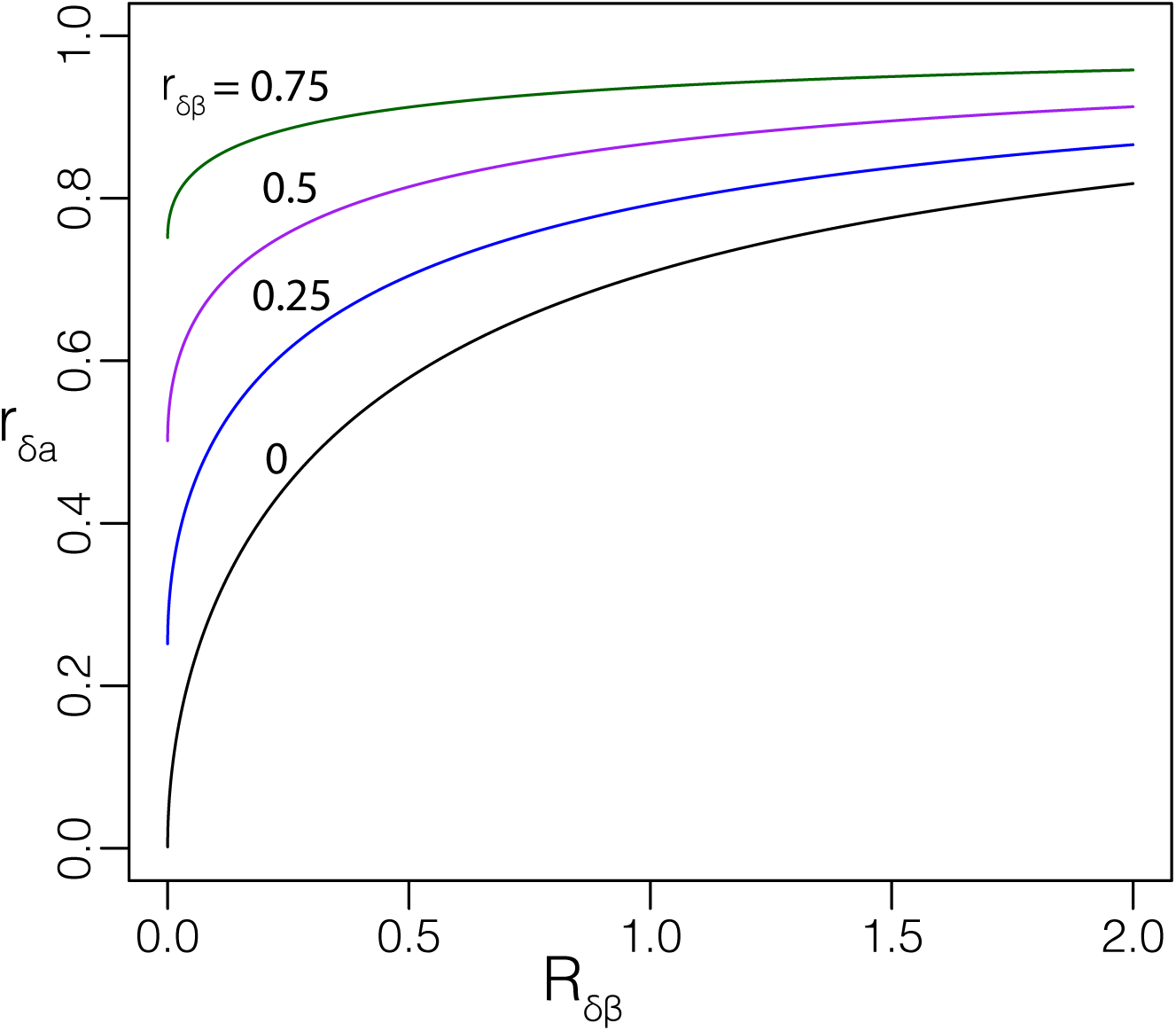
The correlation between direct and population effects. The correlation between direct and population effects (estimated from GWAS on unrelated individuals), *r*_*δa*_, can be expressed as a function of 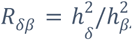, the ratio between the phenotypic variance explained by direct effects and by parental effects, and *r*_*δβ*_, the correlation between direct and parental effects (see above). Here we plot *r*_*δa*_ as a function of *R*_*δβ*_ for various values of *r*_*δβ*_.

We estimated the gain in effective sample from using parental genotypes imputed from sibling genotypes and un-phased IBD information (Figure 5). In the families with at least two genotyped siblings but no genotyped parents, we estimated direct genetic effects using genetic differences between siblings (Methods). In the same sample of families, we compared the standard errors for direct genetic effect estimates from using genetic differences between siblings and from using our method. The gain in effective sample size depends upon both allele frequency and correlation between sibling residuals (Supplementary Note and Figure 2), so varies from SNP to SNP and from trait to trait. For example, the median gain for SNPs with MAF between 1% and 2% was 25.5% for neuroticism, where the sibling correlation is 0.14; whereas the median gain for SNPs with MAF between 49% and 50% was 6.8% for height, where the sibling correlation is 0.52.

**Figure 5.**
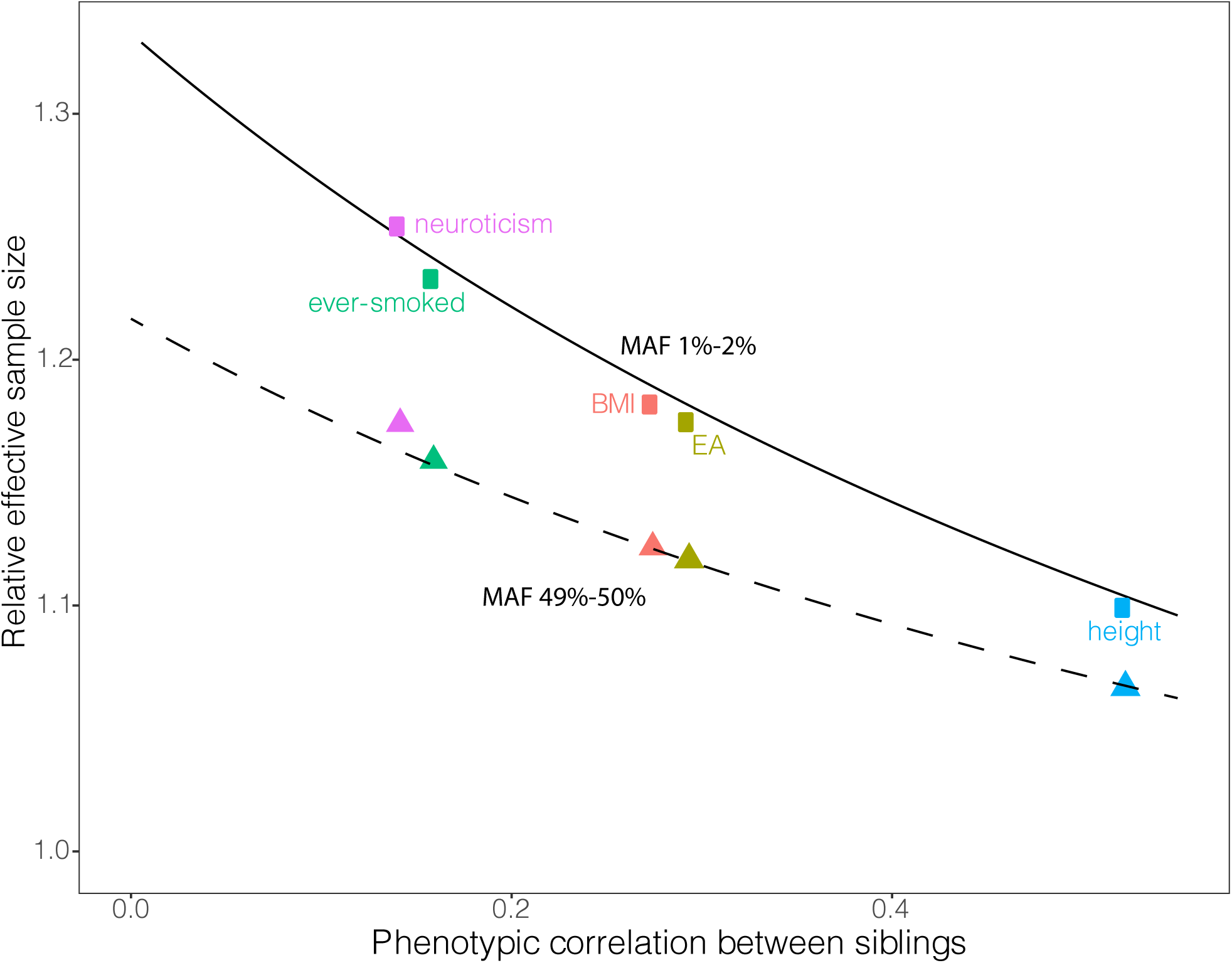
Relative effective sample size for estimation of direct genetic effects for five traits. We show the effective sample size for estimation of direct genetic effects when using parental genotypes imputed from sibling genotypes and un-phased identity-by-descent (IBD) information relative to the effective sample size when using genetic differences between siblings alone (Methods). We show the relative effective sample size for SNPs with minor allele frequency (MAF) between 1% and 2% (squares), and for SNPs with MAF between 49% and 50% (triangles). As expected from theory (Supplementary Note and Figure 2), the relative effective sample size diminishes with increasing phenotypic correlation between siblings. For SNPs with MAF between 1% and 2%, the gain in effective sample size is slightly below the gain that would be expected given access to phased IBD data (Supplementary Note), shown by the solid black curve. For SNPs with MAF between 49% and 50%, the relative effective sample size as a function of the phenotypic correlation between siblings fits well our theoretical result for a SNP with MAF 49.5% (dashed black curve). The gap between the points for MAF 1-2% and MAF 49-50% is due to increased heterozygosity in the more common SNPs, which results in ambiguity in imputation without access to phased IBD data. Trait Abbreviations: BMI, body mass index; EA, educational attainment (years).

## Discussion

Many datasets contain genotype information on siblings but not on the parents of the siblings, and many studies have used genetic differences between siblings to estimate direct genetic effects and to remove bias due to population stratification^7,16,23^. However, these methods do not use all of the information in sibling genotypes. We have shown that, by imputing parental genotypes using IBD information, one can substantially increase the information available for estimation of direct and indirect genetic effects (Figure 2 and Supplementary Note), and one can separately identify direct genetic effects and indirect effects from siblings. While it is possible to impute parental genotypes without using IBD information^18^, we showed that this gains little compared to using IBD information in nearly all scenarios. We also developed a method for imputing the genotype of the missing parent from a parent-offspring pair and showed that one can thereby obtain unbiased estimates of direct genetic effects (Supplementary Note).

We imputed missing parental genotypes from siblings and un-phased IBD information and from parent-offspring pairs in a sample of 45,826 White British individuals from the UK Biobank. Using the imputed parental genotypes, we estimated direct and indirect genetic effects of ∼5 million SNPs on educational attainment, BMI, height, neuroticism, and whether an individual has ever smoked. We did not use phased IBD information due to computational costs of phasing millions of SNPs. However, the methods and software we developed could be easily adapted to phased data, thereby gaining additional information for estimation of direct and indirect genetic effects (Figure 2 and Supplementary Note). Furthermore, our method can be applied to analyse polygenic scores, as we show in our companion paper^17^.

Population effects, estimated from GWAS in unrelated individuals, capture direct effects and indirect effects from relatives, population stratification effects, and magnification of effects due to assortative mating^3^. While recent studies have indicated that population stratification has affected GWAS studies even after correction for genetic principal components^7,10^, questions remain about the overall magnitude of bias. One way to quantify how large the bias in population effect estimates is relative to the signal from direct genetic effects is through measuring the correlation between direct and population effects.

Our results show that population effects are biased estimates of direct effects for all five traits analysed. Any bias in effects on height is likely to be small, but the bias could be substantial for EA, BMI, neuroticism, and whether an individual has ever smoked. It is unlikely that indirect genetic effects alone could explain this for a trait like EA, where the parental phenotype that affects offspring EA is likely to be substantially genetically correlated with offspring EA, leading to correlation between direct and parental effects. Population stratification effects are more likely to be uncorrelated with direct effects, so are a more likely explanation for these results. Nevertheless, our results are consistent with near zero correlations between direct and non-direct components, along with substantial non-direct components (Figure 4 and Supplementary Table 4). Further studies including large numbers of first-degree relative pairs will be needed to see if this result holds in other datasets and to get more precise estimates. Larger sample sizes would also enable decomposition of genetic relations between traits into direct and non-direct components^3^.

If population effects are not highly correlated with direct effects, this has important implications for the potential of genetic prediction in different scenarios. For example, prediction of trait differences between embryos relies on genetic variation within a family, where only direct genetic effects are relevant. If direct genetic effects are not highly correlated with population effect estimates, then embryo selection based on population effect estimates will perform poorly relative to using direct effects^24^. Other implications may depend upon the source of the bias in population effect estimates. If it is primarily population stratification, then this may affect portability of genetic prediction both across populations^25,26^ and within populations^16^. If it is primarily indirect genetic effects, then this implies that substantial gains in predictive ability could be obtained from models that incorporate direct and indirect genetic effects along with genotypes of parents and siblings.

Collection of genetic data on large numbers of families is inevitable as sample sizes grow larger. However, the size of these samples is dwarfed by the samples of distantly related individuals. We see individual level data as one possible pattern of missing data in a vision for human genetic analysis that treats the nuclear family as the fundamental unit of analysis rather than the individual^17^. Individual level data, along with other missing data patterns, can be used to increase the precision of estimates of direct and indirect/parental effects using a form of multivariate meta-analysis^17^ (Supplementary Note). We see such methods as the start of a suite of methods that can powerfully analyse and combine information from different patterns of observed genetic data from families to build a richer and more accurate picture of the role of genetics in human trait variation.

## Acknowledgements

The study was supported by funding from the Li Ka Shing Foundation, the Ragnar Söderberg Foundation (E42/15), Open Philanthropy (010623-00001), and the NIA/NIH through grants R24-AG065184 to the University of Southern California, K99-AG062787-01 to Massachusetts General Hospital, and R21AG067585 to the Broad Institute. We thank the UK Biobank (application ID 11867).

## Software

The software used in this paper is available as a *Python* package with command line tools at https://github.com/AlexTISYoung/SNIPar with documentation at https://sibreg.readthedocs.io/en/master/. We recommend reading the guide (https://sibreg.readthedocs.io/en/master/guide.html) and working through the tutorial (https://sibreg.readthedocs.io/en/master/tutorial.html).

The code is written in C/Python and is multi-threaded. To give a sense of runtime, we give runtimes for analysing the genotyping array SNPs on chromosome 1 in the UK Biobank using a single thread for computation. To impute all of the SNPs on chromosome 1 on the UK Biobank array (∼58,000) for around 20,000 families took around 40 minutes. To estimate the effects for ∼50,000 SNPs and a sample of around ∼40,000 individuals, it took around 18 minutes. These runtimes could be reduced substantially by increasing the number of threads.

## Data

Full summary statistics for direct and indirect effects will be released on publication.

## Methods

### UK Biobank sample

We used the UK Biobank sample that had been identified by UK Biobank to have predominantly White British ancestry^19^. We filtered out individuals identified by UK Biobank to have excess relatives, excess heterozygosity, or sex chromosome aneuploidy. We used the kinship coefficients computed by UK Biobank to identify individuals with a first degree relative, where a first degree relation is defined as a kinship coefficient of 0.177 and above^20^.

We extracted the genotypes for that subsample of the UK Biobank, removing SNPs with missingness above 5%. We used KING^20^ with the ‘--related --degree 1’ options to infer the sibling and parent-offspring relations within that set of individuals. We identified 157 duplicates/monozygotic twins and removed one from each pair from further analyses. We identified 17,296 families with at least two siblings, giving a total of 19,329 sibling pairs. The maximum number of siblings in a family was 6, and 913 families had more than two siblings. We used age and sex information to determine the father/mother in each inferred parent-offspring relation, requiring parents to have a reported age at least 12 years higher than their inferred child; parent-offspring relations with a lower reported age difference were removed from further analyses. We identified 4,418 families with at least one parent and one child genotyped; 736 families had at least one child and the father genotyped but not the mother genotyped; 2,798 families had at least one child and the mother but not the father genotyped; 893 families had at least one child and both parents genotyped. We identified 31 families with two children and both parents genotyped, ‘quads’.

### UK Biobank phenotypes

We performed family based GWAS on educational attainment (EA); standing height (Data Field 50); body mass index (BMI) (Data Field 21001); neuroticism score (Data Field 20127); and whether an individual answered that they had ever smoked or not (Data Field 20160), encoded as a binary variable. For EA, we converted the answers to the qualifications question (Data Field 6138) to years of education according to the method used in previous GWAS of EA^23^. For all traits, we regressed out age, age^2^, age^3^, sex, and interactions between sex and age, age^2^, age^3^, along with the 40 genetic principal components provided by UK Biobank. For quantitative traits measured on a continuous scale (height and BMI), we performed an inverse normal transformation on the residuals separately for males and females and then combined the male and female samples.

### IBD inference and imputation in UK Biobank

We inferred IBD segments between all first degree relatives using the KING --ibdsegs option. We confirmed the accuracy of the IBD segment inference by using the 31 white British families where two siblings and both of their parents have been genotyped. When both parents are heterozygous, the IBD state of the siblings is equal to 2 minus the absolute difference in the siblings’ genotypes, except when both siblings are heterozygous (Supplementary Note). We smoothed the true IBD inferred from the quads to account for genotyping errors: if the IBD state at a SNP differed from its two immediately adjacent neighbours, and both adjacent neighbours had the same IBD state, we changed the IBD state of the SNP to be the same as its neighbours. We computed the fraction of sites inferred to be IBD 0, 1, and 2 given the true IBD state (Supplementary Table 1). The overall probability of inferring the correct IBD state was estimated to be 98.4%.

We imputed missing parental genotypes for the bi-allelic SNPs with INFO>0.99 and MAF>1%. We used hard-call genotypes with a stringent INFO threshold so that any influence of genotype errors on the imputation procedure would be minimal. We examined the bias in the imputed parental genotypes by performing the imputation for the 31 families with two genotyped siblings and both parents genotyped (ignoring the parental genotypes), allowing us to compare the imputed parental genotypes to the observed parental genotypes. If the imputation is unbiased, then the regression coefficient of the imputed parental genotypes onto the observed parental genotypes should be 1. This is because the covariance between the imputed parental genotypes and the observed parental genotypes should be equal to the variance of the imputed parental genotypes (Supplementary Note).

Using 166,587,490 SNP observations, we estimated the regression coefficient to be 0.996. This shows the imputation from the siblings based upon the imputed genotypes is very close to unbiased genome-wide.

We imputed the missing parent’s genotype from the observed parent and full sibling offspring using the procedure outlined above and in the Supplementary Note. To check the imputation, we set one parent missing from the 893 families with both parents genotyped, and we imputed the missing parent using the observed parent and offspring genotypes. The coefficient from regression of the observed parental genotype onto the imputed parental genotype was 0.995, indicating the imputation was approximately unbiased.

### Linear Mixed Model

Phenotype observations from the same family are correlated through both shared genetic factors and shared environmental factors. In order to obtain efficient estimates of SNP effects from phenotypic observations from multiple members of the same family, the phenotypic correlations between members of the same family should be modelled. One way to do this is within a linear mixed model where the mean phenotype within each family is modelled as a random effect. Let *Y*_*i*;_ be the mean-centred phenotype of individual *j* in family *i*, then

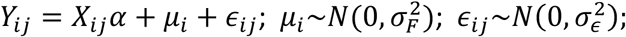

where *X*_*i*j_ are the mean-centered observed/imputed genotypes of individual *j* in family *i*; *α* is the vector of effects; *μ*_*i*_ is the mean in family *i*, which we model as a mean-zero normally distributed random effect with variance 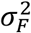, independent for each family; and *ϵ*_*i*j_ is the residual for individual *j* in family *i*, independent for each individual. This implies that, conditional on *X*, the correlation between individuals in the same family is 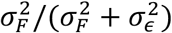.

For estimation of the effects of genome-wide SNPs, we first infer the variance components 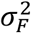 and 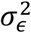 by maximum likelihood for a null model without any SNP effects (Supplementary Note). We then fix the variance components at their maximum likelihood estimate for estimation of the SNP effects. Given the variance components, the maximum likelihood estimate of the vector of effects for a SNP can be obtained analytically in O(n) computations by summing over n families (Supplementary Note). Our software package, SNIPar, performs genome-wide estimation of direct and indirect effects from observed proband and sibling genotypes and observed/imputed parental genotypes.

### Estimation of Effects

We estimated effects for all variants with INFO>0.99 and MAF>1% using the above linear mixed model. Note that although ‘ever-smoked’ was a binary variable, we used a linear model, as in previous GWAS of smoking behaviour^27^. To enable estimation under different models from one analysis of the data, for each SNP, we formed summary statistics from fitting the linear mixed model corresponding to the *X*^T^ *X* matrix and *X*^T^ *Y* vector in standard multivariate linear regression. For the subsample of families with at least two genotyped siblings and no parents genotyped, the *X* matrix had columns corresponding to the proband’s genotype, the mean genotype of the proband’s siblings, and the imputed parental genotype. Let *Y* be the vector of phenotype observations and let Σ = V*a*r(*Y*) be the phenotypic covariance matrix, then the estimate of the parameters under the full model with direct, sibling, and parental effects is the solution to the linear system:

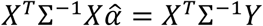

The estimate under the model without sibling effects is obtained by dropping the rows and columns corresponding to the genotype of the proband’s sibling from *X*^T^ Σ^−1^*X* and *X*^T^ Σ^−1^*Y*. Standard GWAS estimates are obtained by using only the rows and columns corresponding to the proband genotype. Note that this method assumes that the variance contribution of each SNP is a very small fraction of the phenotypic variance, so that the residual variance changes only negligibly when certain effects are dropped from the model for each SNP.

For the subsample of families with one parent genotyped, the *X* matrix has columns corresponding to proband genotype, (imputed) paternal, and (imputed) maternal genotypes. For the subsample with both parents genotyped, the *X* matrix has columns corresponding to proband genotype, paternal genotype, and maternal genotype. We did not fit indirect genetic effects from siblings for these subsets of families because only a small fraction of these families had more than one genotyped sibling.

Direct effect estimates from the different subsamples were combined using fixed effects meta-analysis. Indirect sibling effects were estimated from the subsample of families with at least two siblings genotyped and no parents genotyped. For parental effects, we used the multivariate meta-analysis method outline in the Supplementary Note to get meta-analysis estimates of maternal and paternal effects separately, and we took the average of those estimates to give meta-analysis estimates of the average parental effect.

For the subset of families with at least two siblings genotyped but no parents genotyped, we also implemented the difference in sibling genotypes method. We computed the mean genotype of the siblings in each family 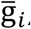, and fit the linear mixed model:

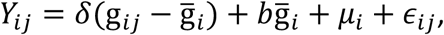

where *b* captures both direct and parental effects.

### LD Score Regression Analysis

To apply LD-score regression to direct, indirect, and population effects, we adjusted the sample size input to LD-score regression to reflect the effective sample size for each effect at each SNP. Note that the effective sample size is considerably smaller for estimation of direct and indirect effects than for population effects. Let 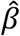 be the effect estimate for a SNP with allele frequency *f* and with sampling variance V*a*r 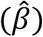. We estimated the effective sample size, *N*_eff_, to be

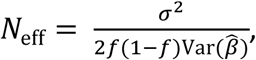

where *σ*^2^; is the phenotypic variance.

For the genetic correlation analysis, we used direct effects and parental effects estimated assuming that indirect effects from siblings are absent.

## Notes

https://github.com/AlexTISYoung/SNIPar

### Competing Interest Statement

The authors have declared no competing interest.

