## Supplementary Information for "Mendelian imputation of parental genotypes for genome-wide estimation of direct and indirect genetic effects"

### 1 Supplementary Figures

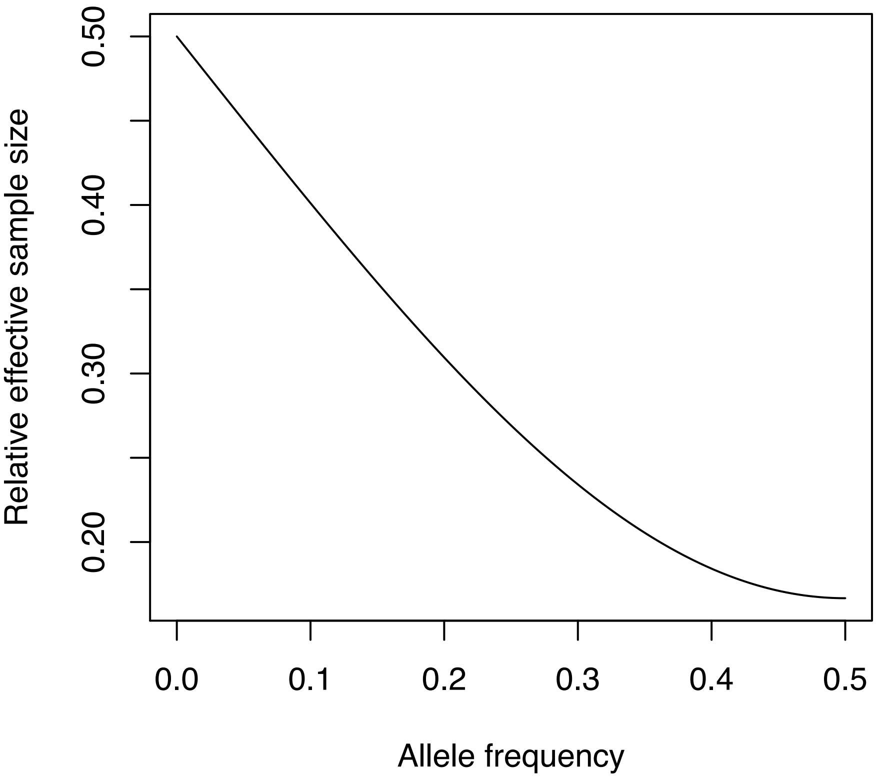

Figure 1: Relative effective sample size for estimating direct genetic effects using paternal genotype imputed from maternal and proband genotype compared to observed paternal genotype as a function of allele frequency.

### 1 Supplementary Tables

| | | Inferred IBD | | |
|---|---|---|---|---|
| | | 0 | 1 | 2 |
| | 0 | 0.997 | 0.002 | 0.000 |
| True IBD | 1 | 0.017 | 0.982 | 0.001 |
| | 2 | 0.002 | 0.023 | 0.975 |

Table 1: The estimated probability of each inferred IBD state given the true IBD state, rounded to 3 decimal places. True IBD states were inferred from 31 families with two siblings and both parents genotyped (Methods and Supplementary Note).

| | | direct | | | direct and parental | |
|---|---|---|---|---|---|---|
| parameter | True | Est. | S.E. | True | Est. | S.E. |
| $h^2_\delta$ | 0.4 | 0.470 | 0.044 | 0.144 | 0.167 | 0.005 |
| $h^2_\beta$ | 0 | - | - | 0.144 | 0.225 | 0.005 |
| $h^2_a$ | 0.4 | 0.432 | 0.024 | 0.432 | 0.363 | 0.003 |
| $r_{\delta\beta}$ | - | - | - | 0.5 | 0.497 | 0.028 |
| $r_{\delta a}$ | 1 | 1.034 | 0.022 | 0.866 | 0.888 | 0.004 |

Table 2: LD score regression results for simulated traits. We give estimates of the variance components $h^2_\delta$, the variance explained by direct effects; $h^2_\beta$, the variance explained by indirect effects from parents; $h^2_a$, the variance explained by the 'population effects', $(\delta + \beta)$; $r_{\delta\beta}$, the correlation between direct and parental effects; and $r_{\delta a}$, the correlation between $\delta$ and the population effects. We simulated one replicate of a trait affected by only direct genetic effects and Gaussian noise, with the direct effects explaining 40% of the phenotypic variance (Methods), called the 'direct' trait. We simulated 40 replicates of a trait affected by both direct and parental effects, where the direct and parental effects each explained 14.4% of the phenotypic variance and were correlated 0.5 ($r_{\delta\eta}$), giving a correlation between direct and population effects of 0.866 ($r_{\delta a}$).

| trait | $h^2_{\text{sib}}$ | S.E. |
|---|---|---|
| EA | -0.0478 | 0.0756 |
| height | 0.0341 | 0.0845 |
| BMI | 0.033 | 0.0857 |
| neuroticism | 0.0719 | 0.085 |
| ever-smoked | -0.1148 | 0.0623 |

Table 3: Estimate of the phenotypic variance explained by indirect effects from siblings from LD-score regression along with their standard errors.

| trait | $r_{\delta\beta}$ | S.E. | $r_{\delta a}$ | S.E. | $P(r_{\delta a} < 1)$ |
|---|---|---|---|---|---|
| EA | -0.3059 | 0.1974 | 0.6110 | 0.0922 | $1.2 \times 10^{-5}$ |
| height | 0.4189 | 0.2571 | 0.9588 | 0.0148 | $2.7 \times 10^{-3}$ |
| BMI | -0.2545 | 0.1635 | 0.8710 | 0.0363 | $1.9 \times 10^{-4}$ |
| neuroticism | -0.5796 | 0.2320 | 0.6793 | 0.1030 | $9.2 \times 10^{-4}$ |
| ever-smoked | -0.7823 | 0.1338 | 0.7274 | 0.0926 | $1.6 \times 10^{-3}$ |

Table 4: LD-score regression estimates of the genetic correlation between direct and average parental effects, $r_{\delta\beta}$, and the genetic correlation between direct and population effects, $r_{\delta a}$. The estimates are given along with their standard errors. The right-most column gives the P-value for a one-sided Z-test for $r_{\delta a} < 1$. Estimates are derived from 45,826 White British individuals in the UK Biobank who have at least one genotyped first degree relative, where missing parental genotypes have been imputed using the methods outlined in the Main Text, Methods and Supplementary Note. To maximise precision, estimates of direct and parental effects were derived assuming that indirect effects from siblings are zero (Supplementary Note).

| trait | study | $r_{\delta a}$ | S.E. | $r_{aa}$ | S.E. |
|---|---|---|---|---|---|
| EA | Lee et al. 2018[1] | 0.5865 | 0.114 | 0.9938 | 0.0493 |
| height | Yengo et al 2018[2] | 1.0148 | 0.0345 | 1.0047 | 0.0151 |
| BMI | Yengo et al. 2018[2] | 0.9583 | 0.0667 | 0.9533 | 0.0221 |
| neuroticism | Nagel et al. 2018[3] | 0.9265 | 0.1892 | 1.0331 | 0.077 |
| ever-smoked | Liu et al. 2019[4] | 0.6367 | 0.0921 | 0.9265 | 0.0808 |

Table 5: LD-score regression estimates of the genetic correlation between direct effects and GWAS effects from publicly available summary statistics, $r_{\delta a}$, and the genetic correlation between GWAS effects from our study and from publicly available summary statistics, $r_{aa}$. The estimates are given along with their standard errors. Our estimates are derived from 45,826 White British individuals in the UK Biobank who have at least one genotyped first degree relative, where missing parental genotypes have been imputed using the methods outlined in the Main Text, Methods and Supplementary Note. To maximise precision, estimates of direct and parental effects were derived assuming that indirect effects from siblings are zero (Supplementary Note). We note that the definition of the smoking trait in Liu et al.[4] is slightly different to ours, potentially explaining why the estimate of $r_{aa}$ is below 1.

### Supplementary Note for: 'Mendelian imputation of parental genotypes for genome-wide estimation of direct and indirect genetic effects'

June 30, 2020

Alexander I. Young

**7 Multivariate Meta-analysis** **17**

**A Conditional Independence Lemma** **18**

### 1 Nuclear family model

Direct genetic effects are effects of alleles in an individual on that individual's trait. Indirect genetic effects are the effects of alleles in one individual on another individual's trait.

Consider a sample of $n$ independent families with two siblings in each family. We consider a model for a single SNP with direct effects and indirect effects from parents and a single sibling:

$$Y_{i1} = \delta g_{i1} + \eta_s g_{i2} + \beta_p g_{p(i)} + \beta_m g_{m(i)} + e_{i1}, \tag{1}$$

$$Y_{i2} = \delta g_{i2} + \eta_s g_{i1} + \beta_p g_{p(i)} + \beta_m g_{m(i)} + e_{i2}, \tag{2}$$

where $Y_{ij}$ is the observed phenotype value for sibling $j$ in family $i$, $g_{ij}$ is the allele count for sibling $j$ in family $i$, $g_{p(i)}$ is the allele count for the father in family $i$, $g_{m(i)}$ is the allele count for the mother in family $i$, $\delta$ is the direct effect of the SNP, $\eta_s$ is the indirect genetic effect from the sibling, $\beta_p$ is the indirect genetic effect from the father, $\beta_m$ is the indirect genetic effect from the mother, and $e_{ij}$ is the residual for individual $j$ from family $i$. For convenience, we assume that phenotypes and genotypes have been mean mean-centred.

#### 1.1 Multiple siblings

Families have different numbers of siblings, with different distributions of ages and genders, among other potentially important variables. While the indirect genetic effect from a single sibling may depend upon the number of other siblings and other factors, for simplicity, we model the indirect effect from multiple siblings as an additive effect of the average of the siblings' genotypes:

$$Y_{ij} = \delta g_{ij} + \eta_s \bar{g}_{i,j} + \beta_p g_{p(i)} + \beta_m g_{m(i)} + e_i, \tag{3}$$

where $\bar{g}_{i,j}$ is the average genotype of the siblings of individual $j$ in family $i$.

#### 1.2 Unbiased estimation of direct and sibling effects

The goal of the analysis is to estimate the parameter vector $\theta = [\delta, \eta_s, \eta_p, \eta_m]^T$. The residual $e_i$ can be correlated with the genotypes due to population stratification. We show that this introduces bias to estimates of $\eta_m$ and $\eta_p$ but not $\delta$ and $\eta_s$. The unbiasedness of the estimates of $\delta$ and $\eta_s$ follows from the fact that, given $g_{p(i)}$ and $g_{m(i)}$, $g_{i1}$ and $g_{i2}$ are conditionally independent of $e_{i1}$ and $e_{i2}$. This follows from the fact that, given parental genotypes, offspring genotypes are determined by random Mendelian segregations in the parents, which are independent of environmental effects [1]. Furthermore, the expectations of $g_{i1}$ and $g_{i2}$ given $g_{p(i)}$ and $g_{m(i)}$ are a linear function of $g_{p(i)}$ and $g_{m(i)}$: $(g_{p(i)} + g_{m(i)})/2$. Therefore, by the Conditional Independence Lemma (Appendix A),

$$Y_{i1} = \delta g_{i1} + \eta_s g_{i2} + (\eta_p + b_p) g_{p(i)} + (\eta_m + b_m) g_{m(i)} + \epsilon_{i1}, \tag{4}$$

$$Y_{i2} = \delta g_{i2} + \eta_s g_{i1} + (\eta_p + b_p) g_{p(i)} + (\eta_m + b_m) g_{m(i)} + \epsilon_{i2}, \tag{5}$$

for some constants $b_p$ and $b_m$ and some $\epsilon_{i1}$ such that $\text{Cov}(g_i, \epsilon_{i1}) = \text{Cov}(g_{i2}, \epsilon_{i1}) = \text{Cov}(g_{p(i)}, \epsilon_{i1}) = \text{Cov}(g_{m(i)}, \epsilon_{i1}) = 0$.

Therefore, $\mathbf{Y}_i = X_i(\theta + \mathbf{b}) + \epsilon_i$, where

$$X_i = \begin{bmatrix} g_{i1} & g_{i2} & g_{p(i)} & g_{m(i)} \\ g_{i2} & g_{ii} & g_{p(i)} & g_{m(i)}, \end{bmatrix} \tag{6}$$

$\mathbf{Y}_i = [Y_{i1}, Y_{i2}]^T$, $\epsilon_i = [\epsilon_{i1}, \epsilon_{i2}]^T$,and $\mathbf{b} = [0, 0, b_p, b_m]^T$.

We model the correlations between residuals for siblings, giving the phenotypic covariance matrix for family $i$ with two sibligs as

$$\Sigma_i = \text{Cov}(Y_i) = \sigma_\epsilon^2 \begin{bmatrix} 1 & r \\ r & 1 \end{bmatrix}, \tag{7}$$

where $r = \text{Corr}(\epsilon_{i1}, \epsilon_{i2})$. The generalised least squares estimator for $\theta$ is

$$\hat{\theta} = \left( \sum_{i=1}^{n} X_i^T \Sigma_i^{-1} X_i \right)^{-1} \left( \sum_{i=1}^{n} X_i^T \Sigma_i^{-1} Y_i \right), \tag{8}$$

which has expectation

$$\mathbb{E}[\hat{\theta}] = \theta + \mathbf{b}, \tag{9}$$

implying unbiased estimation of $\delta$ and $\eta_s$.

### 2 Imputation regression theory

In real data, genotypes of one or both parents are often missing. We consider imputing the missing parental genotypes as the conditional expectation of the missing genotype(s) given the observed genotypes. For example, if we have genotypes on two siblings, we can impute the sum of maternal and paternal genotypes as $\hat{g}_{par(i)} = \mathbb{E}[g_{par(i)} | g_{i1}, g_{i2}]$. We first prove that regression using imputation of this kind produces consistent estimates of parameters in general before considering specific applications.

We first prove a lemma that imputation of this kind changes only the variance of the imputed variables in the variance-covariance matrix of the covariates; and that the covariance between the imputed covariates and the true covariates to be imputed is equal to the variance-covariance matrix of the imputed covariates. This then leads to a theorem that least-squares estimation with the resulting imputed covariates gives consistent estimators of the regression coefficients.

**Lemma 1.** Consider two random column vectors $X_0$ and $X_1$ and a statistic $A$, which could be, for example, the IBD state of two siblings at a SNP. Let $\hat{X}_1 = \mathbb{E}[X_1 | X_0, A]$, then $\text{Cov}(X_1, \hat{X}_1) = \text{Var}(\hat{X}_1)$ and $\text{Cov}(X_0, \hat{X}_1) = \text{Cov}(X_0, X_1)$.

*Proof.* First note that $\mathbb{E}[\hat{X}_1] = \mathbb{E}[\mathbb{E}[X_1|X_0, A]] = \mathbb{E}[X_1]$ by the Law of Iterated Expectations. We now compute

$$\text{Cov}(X_1, \hat{X}_1) = \mathbb{E}[X_1 \hat{X}_1^T] - \mathbb{E}[X_1]\mathbb{E}[\hat{X}_1]^T \tag{10}$$

$$= \mathbb{E}[\mathbb{E}[X_1 \hat{X}_1^T | X_0, A]] - \mathbb{E}[\hat{X}_1]\mathbb{E}[\hat{X}_1]^T, \tag{11}$$

by the Law of Iterated Expectations, and using the fact that $\mathbb{E}[\hat{X}_1] = \mathbb{E}[X_1]$. Since conditional on $X_0$ and $A$, $\hat{X}_1$ is constant, we have that $\mathbb{E}[\mathbb{E}[X_1 \hat{X}_1^T | X_0, A]] = \mathbb{E}[\mathbb{E}[X_1|X_0, A]\hat{X}_1^T] = \mathbb{E}[\hat{X}_1 \hat{X}_1^T]$, and therefore

$$\text{Cov}(X_1, \hat{X}_1) = \mathbb{E}[\hat{X}_1 \hat{X}_1^T] - \mathbb{E}[\hat{X}_1]\mathbb{E}[\hat{X}_1]^T = \text{Var}(\hat{X}_1). \tag{12}$$

We use the Law of Total Covariance to compute $\text{Cov}(X_0, X_1)$:

$$\text{Cov}(X_0, X_1) = \mathbb{E}[\text{Cov}(X_0, X_1|X_0, A)] + \text{Cov}(\mathbb{E}[X_0|X_0, A], \mathbb{E}[X_1|X_0, A]). \tag{13}$$

Since $X_0$ is a constant given $X_0$, $\text{Cov}(X_0, X_1|X_0, A) = 0$. Furthermore, $\mathbb{E}[X_0|X_0, A] = X_0$. Therefore,

$$\text{Cov}(X_0, X_1) = \text{Cov}(X_0, \hat{X}_1). \tag{14}$$

□

**Theorem 2.** Let $X = [X_0\ X_1]$; $\hat{X}_1 = \mathbb{E}[X_1|X_0, A]$; $\hat{X} = [X_0\ \hat{X}_1]$; and $Y = X\theta + \epsilon$, where $\epsilon \perp X$. Then $\hat{\theta} = (\hat{X}^T \hat{X})^{-1} \hat{X}^T Y$ is a consistent estimator of $\theta$ provided that $\text{Var}(\hat{X})$ is invertible.

*Proof.* Provided that $\text{Var}(\hat{X})$ is invertible,

$$\lim_{n\to\infty} \hat{\theta} = \text{Var}(\hat{X})^{-1}\text{Cov}(\hat{X}, Y). \tag{15}$$

Using the above Lemma, we have that $\text{Cov}(\hat{X}, Y) = \text{Cov}(\hat{X}, X\theta) = \text{Var}(\hat{X})\theta$. Therefore,

$$\lim_{n\to\infty} \hat{\theta} = \text{Var}(\hat{X})^{-1}\text{Var}(\hat{X})\theta = \theta. \tag{16}$$

□

### 3 Imputation from siblings

Here we outline how to impute parental genotype from siblings without using IBD data, using phased IBD data, and using un-phased IBD data. We give the imputed parental genotypes as a function of the genotypes of a sibling pair and, if applicable, the IBD state of the siblings; and we compute the variance of the resulting imputed parental genotypes, which is used in the next section to compute the sampling variance of the estimates of direct and indirect effects.

#### 3.1 Imputation without IBD

For a family of two siblings, the simplest form of imputation imputes the sum of the parents' genotypes as $\hat{g}_{par(i)} = \mathbb{E}[g_{par(i)}|g_{i1}, g_{i2}]$. The values are[2] :

| | | $g_{i2}$ 0 | 1 | 2 |
|---|---|---|---|---|
| | 0 | $2f/(2-f)$ | $2/(2-f)$ | $2$ |
| $g_{i1}$ | 1 | $2/(2-f)$ | $2+(2f-1)/(1+f(1-f))$ | $2(1+2f)/(1+f)$ |
| | 2 | $2$ | $2(1+2f)/(1+f)$ | $2(1+3f)/(1+f)$ |

Table 1: $\mathbb{E}[g_{par(i)}|g_{i1}, g_{i2}]$.

By using Theorem 2, we have that regression of proband phenotype onto proband genotype and imputed parental genotype gives a consistent estimator $\delta$ provided that $\eta_s = 0$.

##### 3.1.1 Variance of imputed parental genotype

To compute the variance of the estimator, we need to compute the variance of the imputed parental genotype:

$$\text{Var}(\hat{g}_{par(i)}) = \sum_{g_{i1}, g_{i2}} (\hat{g}_{par(i)} - 4f)^2 \mathbb{P}(g_{i1}, g_{i2}) \tag{17}$$

The joint distribution of sibling genotypes can be derived by conditioning on the parental genotypes:

$$\mathbb{P}(g_{i1}, g_{i2}) = \sum_{g_{m(i)}, g_{p(i)}} \mathbb{P}(g_{i1}, g_{i2}|g_{m(i)}, g_{p(i)}) \mathbb{P}(g_{m(i)}, g_{p(i)}). \tag{18}$$

Since sibling genotypes are determined by independent random segregations in the parents, they are conditionally independent given parental genotype. Therefore,

$$\mathbb{P}(g_{i1}, g_{i2}) = \sum_{g_{m(i)}, g_{p(i)}} \mathbb{P}(g_{i1}|g_{m(i)}, g_{p(i)}) \mathbb{P}(g_{i2}|g_{m(i)}, g_{p(i)}) \mathbb{P}(g_{m(i)}, g_{p(i)}). \tag{19}$$

Under assumptions of random mating, the parental genotypes are independent. Therefore,

$$\mathbb{P}(g_{i1}, g_{i2}) = \sum_{g_{m(i)}, g_{p(i)}} \mathbb{P}(g_{i1}|g_{m(i)}, g_{p(i)}) \mathbb{P}(g_{i2}|g_{m(i)}, g_{p(i)}) \mathbb{P}(g_{m(i)}) \mathbb{P}(g_{p(i)}). \tag{20}$$

The above probabilities can be computed (laboriously) by application of Mendelian laws of inheritance and using parental genotype frequencies at Hardy-Weinberg equilibrium.

| | | $g_{i2}$ | | |
|---|---|---|---|---|
| | | 0 | 1 | 2 |
| | 0 | $(1-f)^2(1-f/2)^2$ | $f(1-f)^2(1-f/2)$ | $f^2(1-f)^2/4$ |
| $g_{i1}$ | 1 | $f(1-f)^2(1-f/2)$ | $f(1-f)[1+f(1-f)]$ | $f^2(1-f)(1+f)/2$ |
| | 2 | $f^2(1-f)^2/4$ | $f^2(1-f)(1+f)/2$ | $f^2(1+f)^2/4$ |

Table 2: The joint distribution for two siblings' genotypes: $\mathbb{P}(g_{i1}, g_{i2})$.

The variance of the imputed parental genotypes can be computed from the above joint distribution of sibling genotypes and the corresponding values of the above imputed parental genotypes. We do not give an expression here due to its complexity, but we note that it decreases with increasing heterozygosity.

#### 3.2 Imputation with IBD

Let $g_{par(i)} = g_{m(i)} + g_{p(i)}$ be the sum of the parental genotypes for individual $i$. This can also be written in terms of the parental alleles: $g_{par(i)} = g^p_{m(i)} + g^m_{m(i)} + g^p_{p(i)} + g^m_{p(i)}$, where $g^p_{m(i)}$ is the paternally inherited allele of the mother of $i$, and $g^m_{p(i)}$ is the maternally inherited allele of the father of $i$. For now, we assume that we know which alleles are shared IBD in addition to the overall IBD state (0, 1, or 2), which, for IBD state 1, requires phased data when both siblings are heterozygous. We construct the estimate of $g_{par(i)}$, $\hat{g}_{par(i)}$, to be the expectation of $g_{par(i)}$ given the observed sibling genotypes and the IBD state of the siblings: $\hat{g}_{par(i)} = \mathbb{E}[g_{par(i)} | g_{i1}, g_{i2}, \text{IBD}]$. This gives:

$$\hat{g}_{par(i)} = \begin{cases} g_{i1} + g_{i2} = g_{par(i)}, & \text{if IBD} = 0 \\ g_{i1} + g^k_{i2} + f, & \text{if IBD} = 1 \\ g_{i1} + 2f, & \text{if IBD} = 2, \end{cases} \tag{21}$$

where $k \in \{m, p\}$ is such that $g^k_{i2}$ is not IBD with the alleles inherited by sibling 1 in family $i$.

If we do not have access to phased IBD data, then if both siblings are heterozygous and the IBD state is 1, then the shared allele cannot be determined. In this case, the shared allele is the allele with frequency $f$ with probability $1-f$, and the shared allele is the allele with frequency $1-f$ with probability $f$. This can be derived from considering the relative frequencies of the three observed parental genotypes: when the allele with frequency $f$ is shared, the probability of observing those three parental alleles is $f(1-f)^2$; and when the allele with frequency $(1-f)$ is observed, the probability of observing those three parental alleles is $f^2(1-f)$. Conditional on both siblings being heterozygous and being in IBD state 1, the probability that the allele with frequency $f$ is shared is $f(1-f)^2/[f(1-f)^2+f^2(1-f)] = f(1-f)^2/f(1-f) = 1-f$; this implies that the probability that the allele with frequency $1-f$ is shared is $f$. The imputed parental genotype is therefore the average over these two possibilities:

$$\mathbb{E}[g_{par(i)} | g_{i1} = 1, g_{i2} = 1, \text{IBD} = 1] = f(2+f) + (1-f)(1+f) = 1 + 2f. \tag{22}$$

Let $H_i$ be the event that both siblings are heterozygous, then the imputed parental genotype without phased IBD data is:

$$\hat{g}_{par(i)} = \begin{cases} g_{i1} + g_{i2} = g_{par(i)}, & \text{if IBD} = 0 \\ g_{i1} + g_{i2}^{k} + f, & \text{if IBD} = 1 \text{ and } \neg H_i \\ 1 + 2f, & \text{if IBD} = 1 \text{ and } H_i \\ g_{i1} + 2f, & \text{if IBD} = 2, \end{cases} \tag{23}$$

##### 3.2.1 Multiple siblings

First we prove that, for more than three siblings, it is impossible for all pairs of siblings to be in an IBD 1 state.

Consider three siblings in family $i$. We write the genotype of a sibling in terms of parental alleles as $(g^{j}_{m(i)}, g^{k}_{p(i)})$ for $j, k \in \{m, p\}$. Without loss of generality, consider that the genotype of sibling 1 is $(g^{p}_{m(i)}, g^{p}_{p(i)})$ and that the genotype of sibling 2 is $(g^{p}_{m(i)}, g^{m}_{p(i)})$, so that sibling 1 and 2 are in an IBD 1 state. We now consider if it is possible for a third sibling to be in an IBD 1 state with both sibling 1 and sibling 2.

If sibling 3 inherits the same parental alleles as either sibling 1 or sibling 2, then sibling 3 is in an IBD 2 state with another sibling. There are two more possible inheritance patterns for sibling 3: $(g^{m}_{m(i)}, g^{p}_{p(i)})$, which implies sibling 3 is IBD 0 with sibling 2; and $(g^{m}_{m(i)}, g^{m}_{p(i)})$, which implies sibling 3 is IBD 0 with sibling 2. Therefore, it is impossible for sibling 3 to be IBD 1 with both sibling 1 and sibling 2.

This implies that, to impute parental genotypes with more than two siblings, the problem can be reduced to imputing exactly the parental alleles when at least one sibling pair is in an IBD 0 state; or imputing as if one has observed a single sibling pair in an IBD state 2 if all siblings are in an IBD 2 state with each; or reduced to the problem of imputing from a sibling pair by reducing sets of siblings that are all IBD 2 with each other to a single sibling. (An additional sibling that is in an IBD 2 state with an existing sibling adds no further observed parental alleles.)

##### 3.2.2 Variance of imputed parental genotype

We compute the variance of the parental genotype imputed from a sibling pair. The variance of the imputed parental genotype can be computed by the Law of Total Variance:

$$\text{Var}(\hat{g}_{par(i)}) = \mathbb{E}_{\text{IBD}}[\text{Var}(\hat{g}_{par(i)}|\text{IBD})] + \text{Var}_{\text{IBD}}(\mathbb{E}[\hat{g}_{par(i)}|\text{IBD}]) = \mathbb{E}_{\text{IBD}}[\text{Var}(\hat{g}_{par(i)}|\text{IBD})], \tag{24}$$

since the expectation of the imputed parental genotypes does not depend upon the IBD state of the siblings. When using phased IBD data for imputation, the variance of the imputed parental genotype is directly proportional to the number of observed parental alleles:

$$\text{Var}(\hat{g}_{par(i)}|\text{IBD}) = \begin{cases} 4f(1-f), & \text{if IBD} = 0 \\ 3f(1-f), & \text{if IBD} = 1 \\ 2f(1-f), & \text{if IBD} = 2 \end{cases}. \tag{25}$$

Therefore, since $\mathbb{P}(\text{IBD} = 0) = 0.25$, $\mathbb{P}(\text{IBD} = 1) = 0.5$, and $\mathbb{P}(\text{IBD} = 2) = 0.25$, $\text{Var}(\hat{g}_{par(i)}) = 3f(1-f) = (3/4)\text{Var}(g_{par(i)})$. The imputed parental genotype thus captures three quarters of the variance of the observed parental genotype.

Without phased IBD data, the variation in the parental genotype captured by the imputation is decreased due to the inability to determine the shared allele when both siblings are heterozygous and in IBD state 1. To compute the variance of the imputed parental genotype, we need to compute the variance of the imputed parental genotype given IBD state 1. The imputed parental genotype as a function of the observed sibling genotypes given IBD state 1 is:

| | | $g_{i2}$ | | |
|---|---|---|---|---|
| | | 0 | 1 | 2 |
| | 0 | $f$ | $1+f$ | - |
| $g_{i1}$ | 1 | $1+f$ | $1+2f$ | 2+f |
| | 2 | - | $2+f$ | $3+f$ |

Table 3: $\mathbb{E}[g_{\text{par(i)}}|g_{i1}, g_{i2}, \text{IBD} = 1]$

By considering the probability of observing the three observed parental alleles, one can derive the distribution of the observed sibling genotypes given that the IBD state is 1:

| | | $g_{i2}$ | | |
|---|---|---|---|---|
| | | 0 | 1 | 2 |
| | 0 | $(1-f)^3$ | $f(1-f)^2$ | 0 |
| $g_{i1}$ | 1 | $f(1-f)^2$ | $f(1-f)$ | $f^2(1-f)$ |
| | 2 | 0 | $f^2(1-f)$ | $f^3$. |

Table 4: $\mathbb{P}(g_{i1}, g_{i2}|\text{IBD} = 1)$

From these two tables, one can compute that

$$\text{Var}(\hat{g}_{\text{par(i)}}|\text{IBD} = 1) = [3 - f(1-f)]f(1-f); \tag{26}$$

and therefore

$$\text{Var}(\hat{g}_{\text{par(i)}}) = [3 - f(1-f)/2]f(1-f). \tag{27}$$

This shows that the fraction of the variance in the parental genotype captured by imputation with un-phased IBD data is:

$$\frac{\text{Var}(\hat{g}_{\text{par(i)}})}{\text{Var}(g_{\text{par(i)}})} = \frac{3 - f(1-f)/2}{4}, \tag{28}$$

which decreases with increasing heterozygosity.

### 4 Estimating effects from siblings

For the analyses in this section, we assume that $\eta_s = 0$. This simplifies the theoretical derivations. We note that estimators of direct and parental effects are biased in proportion to $\eta_s$ when $\eta_s \neq 0$ and imputed parental genotypes are used, unless proband, sibling, and imputed parental genotypes are used jointly as covariates in the regression [ref][kong].

The model for the two siblings' phenotypes is:

$$Y_{i1} = \delta g_{i1} + \beta g_{\text{par(i)}} + \epsilon_{i1}; \tag{29}$$

$$Y_{i2} = \delta g_{i2} + \beta g_{\text{par(i)}} + \epsilon_{i2}. \tag{30}$$

Here $\beta$ is the average of the indirect effects from the mother and the father, plus a bias term: $2\beta = \beta_p + \beta_m + b_p + b_m$, where $b_p + b_m$ is the bias term arising from population stratification and assortative mating (see Equation 4).

#### 4.1 Without imputation

The phenotypes of the two siblings can be transformed into two orthogonal variables:

$$Y_{i1} - Y_{i2} = \delta(g_{i1} - g_{i2}) + \epsilon_{i1} - \epsilon_{i2}; \tag{31}$$

$$Y_{i1} + Y_{i2} = \delta(g_{i1} + g_{i2}) + 2\beta g_{\text{par(i)}} + \epsilon_{i1} + \epsilon_{i2}. \tag{32}$$

The first variable, $Y_{i1} - Y_{i2}$, gives information on $\delta$. The second variable gives information on a linear combination of $\delta$ and $\beta$ that, when combined with the information on $\delta$ from the difference in phenotypes, can give an estimate of $\beta$.

By performing regression of differences between siblings' phenotypes onto differences in genotypes, one can estimate $\delta$. Let $\hat{\delta}_\Delta$ be the resulting estimator. It can be shown that $\mathbb{E}[\hat{\delta}_\Delta] = \delta$ when $\eta_s = 0$; and

$$\text{Var}(\hat{\delta}_\Delta) = \frac{(1-r)\sigma_\epsilon^2}{nf(1-f)}. \tag{33}$$

By performing the regression $(Y_{i1} + Y_{i2}) \sim (g_{i1} + g_{i2})$ one obtains an estimate of $\delta + (4/3)\beta$. Let this estimate be $\hat{a}$. It is trivial to show that $\text{Var}(\hat{a}) = (1+r)\sigma_\epsilon^2/(3nf(1-f))$. We can obtain an estimate of $\beta$ as $\hat{\beta}_\Delta = (3/4)(\hat{a} - \hat{\delta}_\Delta)$, with $\mathbb{E}[\hat{\beta}_\Delta] = \beta$. From this, we have that $\text{Var}(\hat{\beta}_\Delta) = 3\sigma_\epsilon^2(2-r)/(8nf(1-f))$. We also have that $\text{Cov}(\hat{\beta}_\Delta, \hat{\delta}_\Delta) = -3\text{Var}(\hat{\delta}_\Delta)/4 = -3(1-r)\sigma_\epsilon^2/(4nf(1-f))$.

#### 4.2 With imputation

Let $\hat{g}_{par(i)}$ be the imputed parental genotype from one of the above methods — without IBD, with phased IBD, and with un-phased IBD. From Lemma 1, we have that

$$\begin{bmatrix} \text{Var}(g_{i1}) & \text{Cov}(g_{i1}, g_{i2}) & \text{Cov}(g_{i1}, \hat{g}_{\text{par(i)}}) \\ \text{Cov}(g_{i1}, g_{i2}) & \text{Var}(g_{i2}) & \text{Cov}(g_{i2}, \hat{g}_{\text{par(i)}}) \\ \text{Cov}(g_{i1}, \hat{g}_{\text{par(i)}}) & \text{Cov}(g_{i2}, \hat{g}_{\text{par(i)}}) & \text{Var}(\hat{g}_{\text{par(i)}}) \end{bmatrix} = f(1-f) \begin{bmatrix} 2 & 1 & 2 \\ 1 & 2 & 2 \\ 2 & 2 & 4v \end{bmatrix}, \tag{34}$$

where $v$ is the fraction of the variance in parental genotype captured by the imputation.

We compute the sampling variance of $\hat{\theta}_s$ from $n$ independent families with two siblings in each family, where

$$\hat{X}_i = \begin{bmatrix} g_{i1} & \hat{g}_{\text{par(i)}} \\ g_{i2} & \hat{g}_{\text{par(i)}} \end{bmatrix} \tag{35}$$

is the design matrix for family $i$, with $g_{i1}$ the genotype of sibling 1 in family $i$, $g_{i2}$ the genotype of sibling 2 in family $i$, and $\hat{g}_{\text{par(i)}}$ the imputed parental genotype for family $i$.

The generalised least-squares estimator is:

$$\hat{\theta} = \left( \sum_{i=1}^{n} \hat{X}_i^T \Sigma_i^{-1} \hat{X}_i \right)^{-1} \left( \sum_{i=1}^{n} \hat{X}_i^T \Sigma_i^{-1} Y_i \right). \tag{36}$$

Assuming that $\beta^2$ is negligible compared to the phenotypic variance,

$$\text{Var}(\hat{\theta}) \approx \left( \sum_{i=1}^{n} \hat{X}_i^T \Sigma_i^{-1} \hat{X}_i \right)^{-1}. \tag{37}$$

We have that:

$$X_i^T \Sigma_i^{-1} X_i = \frac{1}{\sigma_\epsilon^2 (1-r^2)} \begin{bmatrix} g_{i1} & g_{i2} \\ \hat{g}_{\text{par(i)}} & \hat{g}_{\text{par(i)}} \end{bmatrix} \begin{bmatrix} 1 & -r \\ -r & 1 \end{bmatrix} \begin{bmatrix} g_{i1} & \hat{g}_{\text{par(i)}} \\ g_{i2} & \hat{g}_{\text{par(i)}} \end{bmatrix} \tag{38}$$

$$= \frac{1}{\sigma_\epsilon^2 (1-r^2)} \begin{bmatrix} g_{i1}^2 - 2r g_{i1} g_{i2} + g_{i2}^2 & \hat{g}_{\text{par(i)}}(g_{i1} - r(g_{i1}+g_{i2}) + g_{i2}) \\ \hat{g}_{\text{par(i)}}(g_{i1} - r(g_{i1}+g_{i2}) + g_{i2}) & 2(1-r)\hat{g}_{\text{par(i)}}^2 \end{bmatrix} \tag{39}$$

As the sample size increases,

$$\sum_{i=1}^{n} \hat{X}_i^T \Sigma_i^{-1} \hat{X}_i \to \frac{2nf(1-f)}{\sigma_\epsilon^2 (1-r^2)} \begin{bmatrix} 2-r & 2(1-r) \\ 2(1-r) & 4v(1-r) \end{bmatrix}. \tag{40}$$

Therefore,

$$\left( \sum_{i=1}^{n} \hat{X}_i^T \Sigma_i^{-1} \hat{X}_i \right)^{-1} \to \frac{\sigma_\epsilon^2 (1+r)}{8[(2-r)v + r - 1]nf(1-f)} \begin{bmatrix} 4v(1-r) & -2(1-r) \\ -2(1-r) & (2-r) \end{bmatrix}. \tag{41}$$

Using the above, we can derive the large-sample variance of the generalised least squares estimator of $\delta$ and $\beta$ using different imputation methods:

| method | $v$ | $\text{Var}(\hat{\delta})^1$ | $\text{Var}(\hat{\beta})^1$ |
|---|---|---|---|
| no imputation | - | $(1-r)$ | $(3/8)(2-r)$ |
| un-phased IBD | $3/4 - f(1-f)/8$ | $\frac{(1-r^2)(3-f(1-f)/2)}{2[2+r-(1-r/2)f(1-f)]}$ | $\frac{(1+r)(2-r)}{2[2+r-(1-r/2)f(1-f)]}$ |
| phased IBD | $3/4$ | $\frac{3(1-r^2)}{2(2+r)}$ | $\frac{(1+r)(2-r)}{2(2+r)}$ |
| complete data | 1 | $\frac{1-r^2}{2}$ | $\frac{(1+r)(2-r)}{8}$ |

Table 5: The sampling variance of estimators of direct and parental effects using no imputation, imputation using un-phased IBD, using phased IBD, and using complete data (both maternal and paternal genotype observed). These are given for a sample of $n$ independent sibling pairs with a correlation of $r$ between their residuals. [1]The sampling variance is given as the factor that multiplies $\sigma_\epsilon^2/(nf(1-f))$.

Following a similar procedure, it can be shown that

$$\sum_{i=1}^{n} \hat{X}_i^T \Sigma_i^{-1} \hat{Y}_i \to \frac{2nf(1-f)}{\sigma_\epsilon^2(1-r^2)} \begin{bmatrix} (2-r)\delta + 2(1-r)\beta \\ 2(1-r)(\delta + 2v\beta) \end{bmatrix}. \tag{42}$$

This allows us to compute the limit of the generalised least squares estimator:

$$\hat{\theta} \to \frac{1}{4(1-r)[(2-r)v + r - 1]} \begin{bmatrix} 4v(1-r) & -2(1-r) \\ -2(1-r) & (2-r) \end{bmatrix} \begin{bmatrix} (2-r)\delta + 2(1-r)\beta \\ 2(1-r)(\delta + 2v\beta) \end{bmatrix} = \begin{bmatrix} \delta \\ \beta \end{bmatrix}. \tag{43}$$

Therefore, the different imputation methods all produce consistent estimators of $\delta$ and $\beta$ given that $\eta_s = 0$.

### 5 One parent missing

#### 5.1 Imputation

We impute the missing parental genotype as the expectation given the observed proband and parent genotypes. Assuming that the father's genotype is missing, this is

$$\hat{g}_{\text{p(i)}} = \mathbb{E}[g_{\text{p(i)}} | g_i, g_{\text{m(i)}}] \tag{44}$$

$$= \frac{2[f(1-f)\mathbb{P}(g_i|g_{\text{m(i)}}, g_{\text{p(i)}} = 1) + f^2\mathbb{P}(g_i|g_{\text{m(i)}}, g_{\text{p(i)}} = 2)]}{(1-f)^2\mathbb{P}(g_i|g_{\text{m(i)}}, g_{\text{p(i)}} = 0) + 2f(1-f)\mathbb{P}(g_i|g_{\text{m(i)}}, g_{\text{p(i)}} = 1) + f^2\mathbb{P}(g_i|g_{\text{m(i)}}, g_{\text{p(i)}} = 2)}, \tag{45}$$

which is derived from application of Bayes' Rule.

By applying the Laws of Mendelian Inheritance to compute the above probabilities, one can derive that:

| | | $g_{\mathrm{m(i)}}$ | | |
|---|---|---|---|---|
| | | 0 | 1 | 2 |
| | 0 | $f$ | $f$ | - |
| $g_i$ | 1 | $1+f$ | $2f$ | f |
| | 2 | - | $1+f$ | $1+f$ |

Table 6: $\mathbb{E}[g_{\mathrm{p(i)}}|g_i, g_{\mathrm{m(i)}}]$

Note that it is impossible for a parent to have two copies of an allele and for the offspring to inherit zero copies, without mutation. (We ignore the possibility of genotyping error here.)

##### 5.1.1 Multiple siblings

We generalise the above approach to perform imputation when multiple sibling offspring of the observed parent are genotyped. Let $n_{ij}$, $j = 0, 1, 2$ be the counts of sibling genotypes equal to 0, 1, and 2 respectively, then, if $g_{m(i)} = 2$, we have

$$\hat{g}_{p(i)} = \begin{cases} \frac{1+2^{n_i-1}f}{1+(2^{n_i-1}-1)f}, & \text{if } n_1 = 0; \\ 1, & \text{if } 0 < n_1 < n_i; \\ \frac{f}{2^{n_i-1}(1-f)+f}, & \text{if } n_1 = n_i. \end{cases} \tag{46}$$

If $g_{m(i)} = 1$, we have

$$\hat{g}_{p(i)} = \begin{cases} \frac{1+2^{n_i-1}f}{1+(2^{n_i-1}-1)f}, & \text{if } n_0 = 0,\ n_2 > 0; \\ 1, & \text{if } n_0 > 0,\ n_2 > 0; \\ \frac{f}{2^{n_i-n_1-1}(1-f)+f}, & \text{if } n_2 = 0,\ n_0 > 0; \\ 2f & \text{if } n_1 = n_i. \end{cases} \tag{47}$$

If $g_{m(i)} = 0$, we have

$$\hat{g}_{p(i)} = \begin{cases} \frac{f}{2^{n_i-1}(1-f)+f}, & \text{if } n_1 = 0; \\ 1, & \text{if } 0 < n_1 < n_i; \\ \frac{1+2^{n_i-1}f}{1+(2^{n_i-1}-1)f}, & \text{if } n_1 = n_i. \end{cases} \tag{48}$$

#### 5.2 Association analysis

Consider a sample of $n$ independent families with one parent and one child genotyped. We assume the genotyped parent is the mother for all families for notational convenience.

The phenotype of the proband from family $i$ can be expressed as

$$Y_i = \delta g_i + \beta_p g_{\text{p(i)}} + \beta_m g_{\text{m(i)}} + \epsilon_i, \tag{49}$$

for some mean-zero $\epsilon_i$ such that $\text{Cov}(g_i, \epsilon_i) = \text{Cov}(g_{\text{p(i)}}, \epsilon_i) = \text{Cov}(g_{\text{m(i)}}, \epsilon_i) = 0$. Note that $\beta_p$ and $\beta_m$ as used here can also include a contribution from indirect effects from siblings, if present.

Let $\hat{X}_p = [\mathbf{g}\,\mathbf{g_m}\,\hat{\mathbf{g}}_\mathbf{p}]$. We consider an estimator formed by regression of $Y$ onto $\hat{X}_p$: $\hat{\theta}_p = (\hat{X}_p^T \hat{X})^{-1} \hat{X}_p^T \mathbf{Y}$. The imputed parental genotype is the conditional expectation given the proband and maternal genotype: $\hat{g}_{\text{p(i)}} = \mathbb{E}[g_{\text{p(i)}} | g_i, g_{\text{m(i)}}]$. This means we can apply Theorem 2 to derive that $\lim_{n\to\infty} \hat{\theta}_p = (\delta, \beta_p, \beta_m)$.

We now derive the sampling variance of $\hat{\theta}_p$: given that $\beta_p^2$ is small relative to the phenotypic variance,

$$\text{Var}(\hat{\theta}_p) \approx \frac{\text{Var}(\hat{X}_p)^{-1}}{n}. \tag{50}$$

To derive $\text{Var}(\hat{g}_{\text{p(i)}})$, we first derive the joint probabilities of the observed genotypes using Bayes' Rule and the Laws of Mendelian Inheritance:

| | | $g_{\text{m(i)}}$ | | |
|---|---|---|---|---|
| | | 0 | 1 | 2 |
| | 0 | $(1-f)^3$ | $f(1-f)^2$ | 0 |
| $g_i$ | 1 | $f(1-f)^2$ | $f(1-f)$ | $f^2(1-f)$ |
| | 2 | 0 | $f^2(1-f)$ | $f^3$ |

Table 7: $\mathbb{P}(g_i, g_{\text{m(i)}})$

From this, we can compute that $\text{Var}(\hat{g}_{\text{p(i)}}) = f(1-f)[1-f(1-f)]$.

By application of Lemma 1, we have that $\text{Cov}(g_i, \hat{g}_{\text{p(i)}}) = \text{Cov}(g_i, g_{\text{p(i)}}) = f(1-f)$, and that $\text{Cov}(g_{\text{m(i)}}, \hat{g}_{\text{p(i)}}) = \text{Cov}(g_{\text{m(i)}}, g_{\text{p(i)}}) = 0$. Therefore,

$$\text{Var}(\hat{X}_p) = \begin{bmatrix} 2 & 1 & 1 \\ 1 & 2 & 0 \\ 1 & 0 & 1-f(1-f) \end{bmatrix}; \tag{51}$$

and therefore

$$\text{Var}(\hat{\theta}_p) \approx \frac{1}{f(1-f)[1-3f(1-f)]} \begin{bmatrix} 2-2f(1-f) & 1-f(1-f) & -2 \\ 1-f(1-f) & 1-2f(1-f) & 1 \\ -2 & 1 & 3 \end{bmatrix}. \tag{52}$$

Let $\hat{\delta}_p$ be the resulting estimator of $\delta$, then

$$\text{Var}(\hat{\delta}_p) \approx \frac{2-2f(1-f)}{[1-3f(1-f)]nf(1-f)} \tag{53}$$

This can be compared to the variance of the estimator of delta with both parental genotypes observed, $\hat{\delta}_{po}$: $\text{Var}(\hat{\delta}_{po}) = (nf(1-f))^{-1}$; and

$$\frac{\text{Var}(\hat{\delta}_{po})}{\text{Var}(\hat{\delta}_{p})} = \frac{1-3f(1-f)}{2-2f(1-f)} \tag{54}$$

The penalty relative to using fully observed parental genotypes increases with the heterozygosity due to the fact that when both observed parent and child genotypes are heterozygous, the allele inherited by the child from the observed parent cannot be determined, so an average over the two possible inheritance patterns is taken as the imputation. This could be overcome by determining the segments of DNA that were inherited by the child from the parent, but that is not always computationally feasible to do for all genotyped/imputed SNPs.

### 6 Mixed model inference

We use a mixed model to account for correlations between individuals within a family. The data are comprised of observations on $n$ families. For family $i$, there are $n_i$ observations, giving a total of $\sum_{i=1}^{n} n_i = N$ observations. We assume that the data has been ordered so that the observations from family 1 are indexed from 1 to $n_1$, the observations from individual 2 are indexed from $n_1 + 1$ to $n_1 + n_2$, etc.

The phenotype is an $[N \times 1]$ vector $Y$ and the covariate matrix is an $[N \times c]$ matrix $X$. The $X$ matrix can be constructed using the genotypes of the siblings and/or (imputed) parents in different ways depending on the application. We introduce a $[N \times n]$ matrix $Z$ such that $[Z]_{ij} = 1$ if observation $i$ is from family $j$, and $[Z]_{ij}$ is zero otherwise. We assume that the within-family means are independently normally distributed, represented by an $[n \times 1]$ vector $u \sim \mathcal{N}(0, \sigma_F^2 \text{I}_n)$. The model is

$$Y = X\alpha + Zu + \epsilon. \tag{55}$$

We assume that the residuals are I.I.D. Gaussians, $\epsilon \sim N(0, \sigma^2 I)$. The distribution of $Y|X$ is therefore,

$$Y|X \sim \mathcal{N}(X\alpha, \sigma_F^2 ZZ^T + \sigma_\epsilon^2 I). \tag{56}$$

It can readily be inferred that $ZZ^T$ has a simple block-diagonal structure. For $n = 2$, the matrix has the following structure:

$$ZZ^T = \begin{bmatrix} 1_{n_1} 1_{n_1}^T & 0 \\ 0 & 1_{n_2} 1_{n_2}^T \end{bmatrix}, \tag{57}$$

where $1_k$ is the $[k \times 1]$ column vector of all 1s.

#### 6.1 Loss function and gradients

Instead of the optimising the likelihood, we seek to minimise negative two times the log-likelihood as a loss function:

$$L = \log|\Sigma| + (y - X\alpha)^T \Sigma^{-1}(y - X\alpha), \tag{58}$$

where $\Sigma = \sigma_F^2 ZZ^T + \sigma_\epsilon^2 I$.

Naive computation of the loss function takes $O(N^3)$ operations. However, the likelihood component of the loss function can be split into a sum over families. Let $\Sigma_i$ be the diagonal block of $\Sigma$ corresponding to observations on family $i$. Furthermore, let $y_i$ be the $[n_i \times 1]$ vector of observations for individual $i$, and let $X_i$ be the $[n_i \times c]$ matrix of covariate observations. Then,

$$L = \sum_{i=1}^{n} \log|\Sigma_i| + \sum_{i=1}^{n} (y_i - X_i\alpha)^T \Sigma_i^{-1}(y_i - X_i\alpha), \tag{59}$$

It is straightforward to show that the MLE for $\alpha$, $\hat{\alpha}$, given the variance parameters, corresponds to the generalised least-squares estimator:

$$\hat{\alpha} = \left(\sum_{i=1}^{n} X_i^T \Sigma_i^{-1} X_i\right)^{-1} \left(\sum_{i=1}^{n} X_i^T \Sigma_i^{-1} y_i\right). \tag{60}$$

It is also straightforward to show that the asymptotic sampling variance of the MLE for $\alpha$ is:

$$\mathrm{Var}(\hat{\alpha}) = \left(\sum_{i=1}^{n} X_i^T \Sigma_i^{-1} X_i\right)^{-1}. \tag{61}$$

To derive the gradient with respect to the variance parameters, we introduce $\tau = \sigma_\epsilon^2/\sigma_F^2$, and parameterise the model in terms of $\tau$ and $\sigma_\epsilon^2$. Because the blocks of $\Sigma$ are comprised of a diagonal plus a rank-one matrix, the determinant and inverse of each block can be computed analytically using the Sherman-Morrison-Woodbury identity and the Matrix Determinant Lemma. This gives

$$\Sigma_i^{-1} = \frac{1}{\sigma_\epsilon^2}\left(\mathrm{I}_{n_i} - \frac{1_{n_i}1_{n_i}^T}{\tau + n_i}\right); \ \log|\Sigma_i| = n_i \log(\sigma_\epsilon^2) + \log\left(1 + \frac{n_i}{\tau}\right). \tag{62}$$

The loss function can thus be expressed as

$$L = N\log(\sigma_\epsilon^2) + \sum_{i=1}^{n} \log\left(1 + \frac{n_i}{\tau}\right) + \frac{(y - X\alpha)^T(y - X\alpha)}{\sigma_\epsilon^2} - \frac{1}{\sigma_\epsilon^2}\sum_{i=1}^{n} \frac{[1_{n_i}^T(y_i - X_i\alpha)]^2}{\tau + n_i}. \tag{63}$$

From this, we have

$$\frac{\partial L}{\partial \sigma_\epsilon^2} = \frac{N}{\sigma_\epsilon^2} - \frac{(y - X\alpha)^T(y - X\alpha)}{\sigma_\epsilon^4} + \frac{1}{\sigma_\epsilon^4}\sum_{i=1}^{n} \frac{[1_{n_i}^T(y_i - X_i\alpha)]^2}{\tau + n_i}; \tag{64}$$

and

$$\frac{\partial L}{\partial \tau} = \frac{1}{\sigma_\epsilon^2}\sum_{i=1}^{n} \frac{[1_{n_i}^T(y_i - X_i\alpha)]^2}{(\tau + n_i)^2} - \sum_{i=1}^{n} \frac{n_i}{\tau(\tau + n_i)}. \tag{65}$$

#### 6.2 Optimisation

The parameters we are optimising over are $\theta = (\alpha, \sigma_\epsilon^2, \tau)$. However, since the MLE for $\alpha$ can be computed efficiently analytically given an estimate of $\tau$, we instead optimise

$$L_{\text{prof}}(\sigma_\epsilon^2, \tau) = L(\hat{\alpha}(\tau), \sigma_\epsilon^2, \tau), \tag{66}$$

where the optimisation takes place over $(\sigma_\epsilon^2, \tau)$ only, with the MLE for $\alpha$ for a given $\tau$, $\hat{\alpha}(\tau)$, computed analytically.

We optimise the model with the L-BFGS-B algorithm, with $(\sigma_\epsilon^2, \tau)$ bounded below at $(10^{-5}, 10^{-5})$. Correct calculation of the loss function was checked by comparing to calculation from the expression in equation 58. Correct calculation of the gradient was checked by numerical differentiation of the loss function. (The checks are performed using unit testing in the *Python* package).

For application to a set of SNPs from a chromosome, a null model including no SNPs is first fit. By default, we initialise $(\sigma_\epsilon^2, \tau)$ to $(s_Y^2/2, 1)$, where $s_Y^2$ is the sample estimate of the phenotypic variance. The MLEs of $\tau$ and $\sigma_\epsilon^2$ from the null model are then fixed for all SNP specific models, allowing analytical computation of the (approximate) MLE for $\alpha$ for each SNP.

### 7 Multivariate Meta-analysis

Consider a parameter vector $\alpha$ and independent observations $z_i \sim \mathcal{N}(A_i\alpha, \Sigma_i)$ for $i = 1, \ldots, k$, then it can be shown that the MLE for $\alpha$ is

$$\hat{\alpha} = \left( \sum_{i=1}^{k} A_i^T \Sigma_i^{-1} A_i \right)^{-1} \left( \sum_{i=1}^{k} A_i^T \Sigma_i^{-1} z_i \right), \tag{67}$$

with $\mathbb{E}[\hat{\alpha}] = \alpha$ and

$$\text{Var}(\hat{\alpha}) = \left( \sum_{i=1}^{k} A_i^T \Sigma_i^{-1} A_i \right)^{-1}. \tag{68}$$

Note that this assumes that $\sum_{i=1}^{k} A_i^T \Sigma_i^{-1} A_i$ is invertible.

Consider estimating parameters in the model

$$Y_i = \delta g_i + \beta_p g_{p(i)} + \beta_m g_{m(i)} + \epsilon_i, \tag{69}$$

where $\epsilon_i$ is uncorrelated with $g_i$, $g_{p(i)}$, and $g_{m(i)}$. We assume that indirect effects from siblings are zero, $\eta_s = 0$. We consider estimating $\theta = [\delta, \beta_p, \beta_m]^T$ using different samples with different observations of sibling and parental alleles. Let sample 1 be a sample where only sibling genotypes are available and the sum of maternal and paternal genotypes have been imputed. Let sample 2 be a sample where proband and maternal genotypes have been observed and paternal genotypes have been imputed . Let sample 3 be a sample where proband, maternal and paternal genotypes have been observed. Then we can combine the estimates from these regressions using the following matrices that give the linear transformation between $\theta$ and the expected estimates from the regressions in each subsample:

| sample | observed genotypes | regression | $\mathbb{E}[z_i]$ | $A_i$ |
|---|---|---|---|---|
| 1 | siblings | $Y_{ij} \sim g_{ij} + \hat{g}_{par(i)}$ | $\begin{bmatrix} \delta \\ 0.5(\beta_p + \beta_m) \end{bmatrix}$ | $\begin{bmatrix} 1 & 0 & 0 \\ 0 & 0.5 & 0.5 \end{bmatrix}$ |
| 2 | proband and maternal | $Y_i \sim g_i + \hat{g}_{p(i)} + g_{m(i)}$ | $\begin{bmatrix} \delta \\ \beta_p \\ \beta_m \end{bmatrix}$ | $I_3$ |
| 2 | proband, paternal, and maternal | $Y_i \sim g_i + \hat{g}_{p(i)} + g_{m(i)}$ | $\begin{bmatrix} \delta \\ \beta_p \\ \beta_m \end{bmatrix}$ | $I_3$ |

Table 8: Linear transformation ($A$) matrices for meta-analysis combining different samples to estimate $\theta = [\delta,\, \beta_p,\, \beta_m]^T$.

### A Conditional Independence Lemma

This Lemma is taken from Young et al. 2018[1], provided here for completeness.

**Lemma 3.** Consider random variables $x$, $y$, and random column vector $z$ such that $x \perp y \,|\, z$, and, for constants $\alpha$ and $b$, and a constant vector of length equal to $z$, $a$,

$$\mathbb{E}[x|z] = \alpha + ba^T z, \tag{70}$$

i.e. the the conditional expectation of $x$ given $z$ is a linear function of some linear combination of the elements of $z$, then

$$\text{Cov}(x, r) = 0, \text{ where } r = y - \frac{\text{Cov}(y, a^T z)}{\text{Var}(a^T z)} a^T z \tag{71}$$

**Remark.** This means that if the expectation of $x$ given $z$ is a linear function of the elements of $z$, and if $x$ is independent of $y$ given $z$, then the residual of the regression of $y$ on $a^T z$ is uncorrelated with $x$. Note that we only assume that there is a linear relationship between $x$ and $z$, not $y$ and $z$.

*Proof.* To prove it, first note that by standard regression theory,

$$b = \frac{\text{Cov}(x, a^T z)}{\text{Var}(a^T z)}, \tag{72}$$

so

$$\text{Cov}(x, r) = \text{Cov}(y, x) - \text{Cov}(y, a^T z) b. \tag{73}$$

It therefore suffices to show that $\text{Cov}(y, x) = \text{Cov}(y, a^T z)b$.

By the Law of Total Covariance

$$\text{Cov}(y, x) = \mathbb{E}_z[\text{Cov}(y, x|z)] + \text{Cov}_z(\mathbb{E}[x|z], \mathbb{E}[y|z]) = \text{Cov}_z(\mathbb{E}[x|z], \mathbb{E}[y|z]), \tag{74}$$

as $\text{Cov}(y, x|z) = 0$, because $x \perp y \mid z$. Therefore,

$$\text{Cov}(y, x) = \text{Cov}_z \left(\alpha + ba^T z, \mathbb{E}[y|z]\right) = \text{Cov}(\mathbb{E}[y|z], a^T z)b \tag{75}$$

It now suffices to show that $\text{Cov}(y, a^T z) = \text{Cov}(\mathbb{E}[y|z], a^T z)$. Without loss of generality, for some $\epsilon$ such that $\mathbb{E}[\epsilon|z] = 0$,

$$y = \mathbb{E}[y|z] + \epsilon. \tag{76}$$

Therefore, $\text{Cov}(y, a^T z) = \text{Cov}(\mathbb{E}[y|z], a^T z) + \text{Cov}(\epsilon, a^T z)$. $\text{Cov}(\epsilon, a^T z) = \mathbb{E}_z[a^T z \mathbb{E}[\epsilon|z]] - \mathbb{E}[a^T z]\mathbb{E}[\epsilon] = -\mathbb{E}[a^T z]\mathbb{E}[\epsilon]$, as $\mathbb{E}[\epsilon|z] = 0$. We also have that $\mathbb{E}[\epsilon] = \mathbb{E}_z[\mathbb{E}[\epsilon|z]] = 0$. Therefore, $\text{Cov}(\epsilon, a^T z) = 0$ and $\text{Cov}(y, a^T z) = \text{Cov}(\mathbb{E}[y|z], a^T z)$, implying

$$\text{Cov}(y, x) = \text{Cov}(y, a^T z)b \Rightarrow \text{Cov}(x, r) = 0. \tag{77}$$

□
